# RSC and GRFs confer promoter directionality by limiting divergent noncoding transcription

**DOI:** 10.1101/2021.08.16.456464

**Authors:** Andrew C.K. Wu, Claudia Vivori, Harshil Patel, Theodora Sideri, Fabien Moretto, Folkert J. van Werven

## Abstract

The directionality of gene promoters - the ratio of protein-coding over divergent noncoding transcription - is highly variable and regulated. How promoter directionality is controlled remains poorly understood. Here, we show that the chromatin remodelling complex RSC and general regulatory factors (GRFs) dictate promoter directionality by attenuating divergent transcription. At gene promoters that are highly directional, depletion of RSC leads to a relative increase in divergent noncoding transcription and thus a decrease in promoter directionality. We find that RSC facilitates nucleosome positioning upstream in promoters at the sites of divergent transcription. These highly directional promoters are also enriched for the binding of GRFs such as Reb1 and Abf1. Ectopic targeting of divergent transcription initiation sites with GRFs or the dCas9 protein suppresses divergent transcription. Our data suggest that RSC and GRFs play a pervasive role in limiting divergent transcription. We propose that any DNA binding factor, when stably associated with cryptic transcription start sites, form barriers for repressing divergent transcription. Our study provides an explanation as to why certain promoters are more directional than others.

## Introduction

Transcription is highly pervasive and results in intergenic and intragenic noncoding transcription events. Noncoding transcription and the produced noncoding RNAs play diverse roles in gene and genome regulation (Ard et al 2017, Gil & Ulitsky 2020).

Transcriptionally active, protein-coding gene promoters are a major source of noncoding transcription. At these genomic locations, noncoding transcription initiates in the divergent direction, a process known as divergent or bidirectional transcription, which generates upstream transcripts from a distinct core promoter in the opposing direction to the coding gene (Neil et al 2009, Seila et al 2008, Sigova et al 2013, Xu et al 2009). Divergent noncoding transcription is an intrinsic property of coding gene transcription, present across eukaryotes, and widespread across actively transcribed regions in the genome (Seila et al 2009). Divergent and coding gene transcription share the same promoter sequence and the ratio (divergent over coding transcription) gives insights on the directionality of promoters (Jin et al 2017).

Our understanding of the function of divergent transcription is still incomplete. There is evidence that the noncoding transcripts emanating from divergent transcription events can help to promote gene transcription in the coding direction and to facilitate cell fate control (Frank et al 2019, Luo et al 2016, Yang et al 2021). Other studies have suggested that divergent transcription may consitute transcriptional noise of active promoters (Seila et al 2009, Struhl 2007). Accordingly, promoters with a high ratio of divergent over coding transcription may have evolved fortuitously, and these promoters have been proposed to represent a transcriptional ground-state (Jin et al 2017, Wu & Sharp 2013). Conversely, highly directional promoters (*i.e.* promoters with little divergent transcription but high levels of sense coding gene transcription) are likely more evolved.

Aberrant noncoding transcription, including divergent transcription, can negatively impact cell fitness. For example, induced noncoding transcription can cause R-loop formation and DNA damage (Nojima et al 2018). In addition, aberrant divergent transcription events can affect coding gene transcription leading to mis-regulation of gene expression (Chiu et al 2018, du Mee et al 2018). This especially impacts species with gene-dense genomes such as *Saccharomyces cerevisiae*. Hence, there are various molecular mechanisms that limit the accumulation of noncoding RNAs or noncoding transcription itself. Indeed, to reduce the accumulation of divergent RNAs, these noncoding transcripts are often rapidly degraded via the RNA degradation machinery (Flynn et al 2011, Malabat et al 2015, Neil et al 2009, van Dijk et al 2011, Xu et al 2009). Divergent transcription units are also typically short, due to enrichment in transcription termination signals (Schulz et al 2013). Additionally, divergent transcription can be repressed by chromatin remodellers and histone modifying enzymes (Gowthaman et al 2021, Marquardt et al 2014). Lastly, general regulatory factors (GRFs) can repress initiation of aberrant noncoding transcription events (Challal et al 2018, Wu et al 2018).

In previous work, we showed that the GRF and pioneer transcription factor Rap1 represses divergent transcription in the promoters of highly expressed ribosomal protein and metabolic genes (Challal et al 2018, Wu et al 2018). At these highly directional promoters, Rap1 is key for both promoting transcription in the coding direction and for limiting transcription in the divergent direction (Challal et al 2018, Wu et al 2018). We proposed that Rap1 limits divergent transcription by interfering with recruitment of the transcription machinery. In the same study, we also identified a role for RSC, a major chromatin remodelling complex important for chromatin organization (Cairns et al 1996). We showed for two gene promoters that RSC promotes divergent transcription in Rap1 depleted cells. RSC activity increases nucleosome positioning, which, in turn, is important for maintaining nucleosome depleted regions (NDRs) in promoters (Badis et al 2008, Hartley & Madhani 2009, Lorch et al 1998, Parnell et al 2008). RSC also acts with GRFs to stimulate transcription in the protein-coding direction (Brahma & Henikoff 2019, Kubik et al 2017, Kubik et al 2018). Specifically, RSC positions the +1 nucleosome in gene promoters to allow pre-initiation complex (PIC) assembly and transcription start site (TSS) scanning (Klein-Brill et al 2019). RSC limits aberrant transcription from upstream in promoters in the sense direction, but also limits transcription initiating from the 3’ end of gene bodies in the antisense direction (Alcid & Tsukiyama 2014, Cucinotta et al 2021, Klein-Brill et al 2019, Kubik et al 2019). The role of RSC in controlling promoter directionality remains an area of interest.

Here, we examined the role of RSC and GRFs in controlling promoter directionality. Our analysis reveals that RSC depletion leads to relative in increase in divergent noncoding transcription. These promoters tend to be highly directional and are enriched for GRFs such as Abf1 and Reb1. Consistent with the role of RSC in chromatin organization, its depletion affects nucleosome positioning in upstream regions of directional promoters. Finally, we demonstrate that ectopic targeting of GRFs and dCas9 to divergent core promoters can also repress divergent noncoding transcription. We propose that nucleosomes positioned by RSC and GRFs constitute physical barriers which limit aberrant divergent transcription in promoters, thereby increasing promoter directionality.

## Results

### RSC represses divergent transcription independently of Rap1

To investigate how RSC and GRFs promote directionality of transcripiton more closely, we performed RNA-seq on nascent transcribed RNA (nascent RNA-seq) and on polyadenylated RNAs (mRNA-seq) after depletion of both RSC and Rap1. We determined the levels of nascently transcribed RNA by measuring RNA Pol II-associated transcripts using an adapted native elongating transcript sequencing protocol (Churchman & Weissman 2011). In short, we affinity-purified RNA Pol II using Rpb3-FLAG and quantified the Pol II-associated RNAs in wild-type (WT) cells, and after depletion of RSC and/or Rap1 (Figure S1A). To deplete RSC and Rap1 we used auxin-inducible degron alleles (*RAP1-AID* and *STH1-AID*) and treated cells with indole-3-acetic acid (IAA) (Nishimura et al 2009). Rap1 and Sth1 were efficiently depleted in cells harbouring either one or both degron alleles after treatment (Figure 1A, top panel). For mRNA-seq we added a spike-in control of *S. pombe* cells in a defined ratio and used the *S. pombe* mRNA-seq signal to normalize mRNA-seq data for *S. cerevisiae* (Figure 1A, bottom panel).

**Figure 1.**
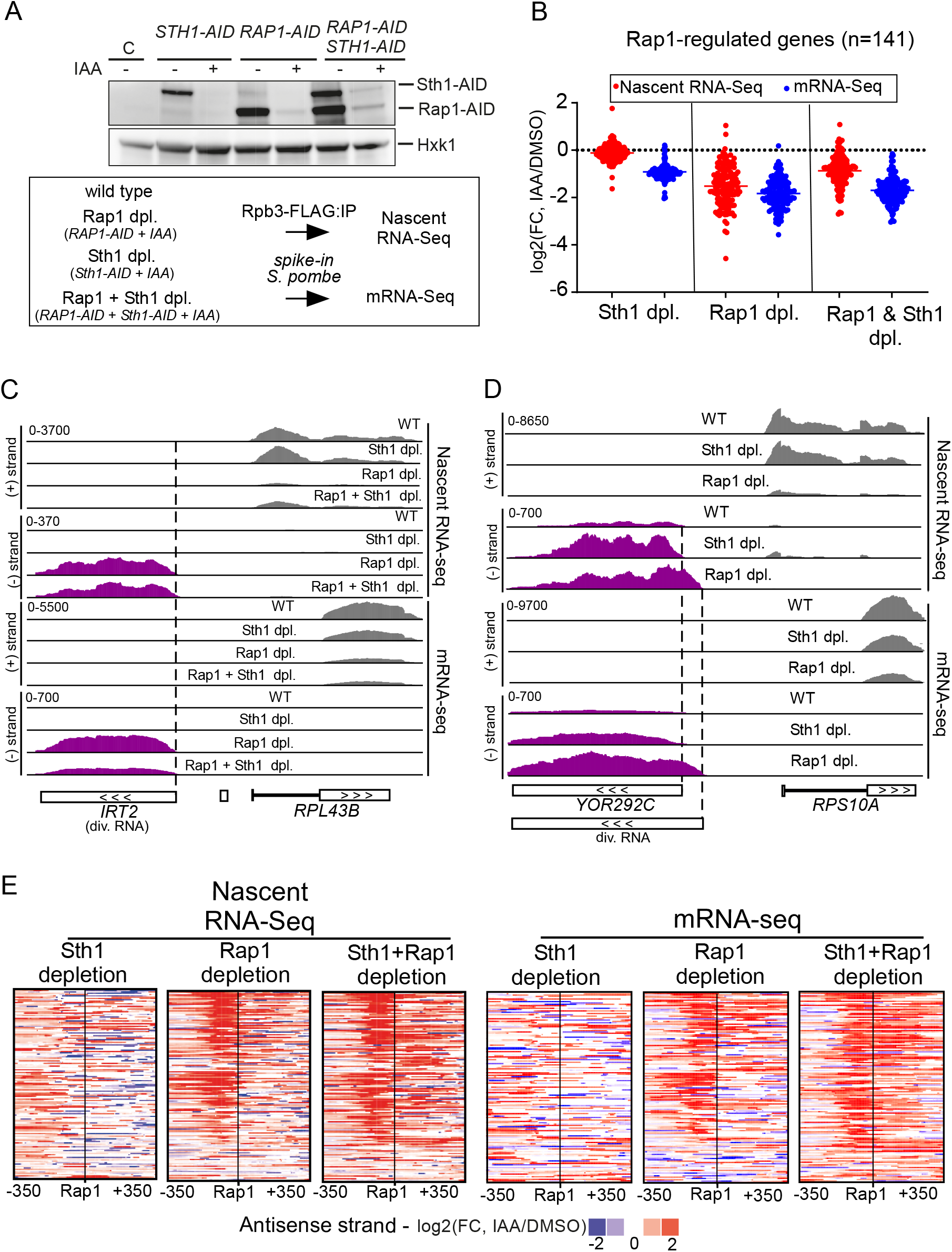
Depletion of RSC increases divergent transcript accumulation at Rap1-regulated genes. **(A)** Top: Rap1 and Sth1 depletions using AID tagged strains (*RAP1-AID*, *STH1-AID*, *RAP1-AID*/*STH1-AID*) (FW7238, FW7220, and FW7232). Cells were grown until the exponential growth phase and treated with IAA for 2 hours. AID tagged proteins were detected by western blot using anti-V5 antibodies. Hxk1 was used as a loading control, detected using anti-Hxk1 antibodies. Bottom: Scheme for nascent RNA-seq and mRNA-seq experiment of Rap1 and Sth1 depletion. To control for global changes in RNA expression, we spiked in *S. pombe* cells (see Materials and Methods), and performed mRNA-seq comparative analysis using *S. pombe* (sp) normalization. **(B)** Fold change (FC) of 141 Rap1-regulated genes nascent RNA-seq and mRNA-seq in IAA treated cells compared to mock treated cells (DMSO). Each dot represents one coding gene transcript of n=3 biological repeats. **(C)** Nascent RNA-seq and mRNA-seq signals at the *RPL43B* locus (coding sense direction, grey) and *IRT2* (divergent coding direction, purple). **(D)** Similar as C except that the *RPS10A* locus and the corresponding divergent transcripts are shown. **(E)** Heatmap representing changes in RNA expression levels in the divergent noncoding direction. Promoters were clustered on antisense strand signal using k-means clustering (k = 3, (c1, c2, and c3)) based on previous analysis for Rap1-regulated gene promoters (Wu et al 2018). Differences with respect to the DMSO control for each depletion strain are displayed.

First, we examined how a subset of regulated protein-coding transcripts were affected by Rap1 and RSC depletion. We focused on a set of 141 Rap1-regulated gene promoters described previously (Wu et al 2018). As expected, almost all Rap1-regulated genes decreased in expression after Rap1 depletion (*RAP1-AID* + IAA vs *RAP1-AID* + DMSO) (Figure 1B). Expression of Rap1-regulated genes also decreased in Sth1-depleted cells as detected by mRNA-seq (Figure 1B). This decrease was lesser in nascent RNA-seq likely, in part, because internal normalization was used for the analysis. Second, we examined two previously well-characterized promoters (*RPL43B* and *RPL40B*) at which Rap1 is known to repress divergent transcription (Figures 1C and S1B) (Wu et al 2018). As expected, transcription of divergent noncoding transcripts (*IRT2* and *iMLP1*) was upregulated upon Rap1 depletion, while coding transcription decreased. Upon co-depletion of Rap1 and Sth1, we observed that *IRT2* and *iMLP1* levels were reduced in mRNA-seq. While Sth1 depletion by itself did not affect *IRT2* levels, levels of *iMLP1* were already upregulated in Sth1-depleted cells (Figure S1B). This suggests that RSC may repress divergent transcription at this locus. Indeed, the *RPS10A* gene showed increased divergent transcription in both Rap1-and Sth1-depleted cells (Figure 1D). Interestingly, the divergent transcription in Sth1-depleted cells initiated further upstream from the coding gene than in Rap1-depleted cells, suggesting that Rap1 and RSC limit divergent transcription through different mechanisms (Figure 1D).

Next, we examined Rap1-regulated gene promoters more closely for changes in divergent transcription (Figure 1E). As expected, Rap1-regulated gene promoters displayed increased levels of divergent transcripts upon Rap1 depletion (Figure 1E). In Sth1-depleted cells we also observed a notable increase in expression of divergent RNAs in nascent RNA-seq (see Figure 2C for quantification) and, to a lesser extent, in the mRNA-seq (Figure 1E). The changes in divergent RNA transcription after Sth1 and Rap1 co-depletion were comparable to Rap1 depletion, suggesting that at Rap1-regulated genes in Rap1 depleted cells RSC does not further promote or repress divergent transcription. Notably, we observed increased RNA expression in the protein-coding direction in Rap1-depleted and Rap1/Sth1 co-depleted cells (Figure S1C). Indeed, it is known that Rap1 depletion can lead to inappropriate transcription initiation in both the protein-coding and divergent directions (Challal et al 2018). These data suggest that RSC depletion caused a relatively greater in divergent transcription compared to protein coding transcription at Rap1 regulated gene promoters, which we further explored in Figures 2 to 5.

**Figure 2.**
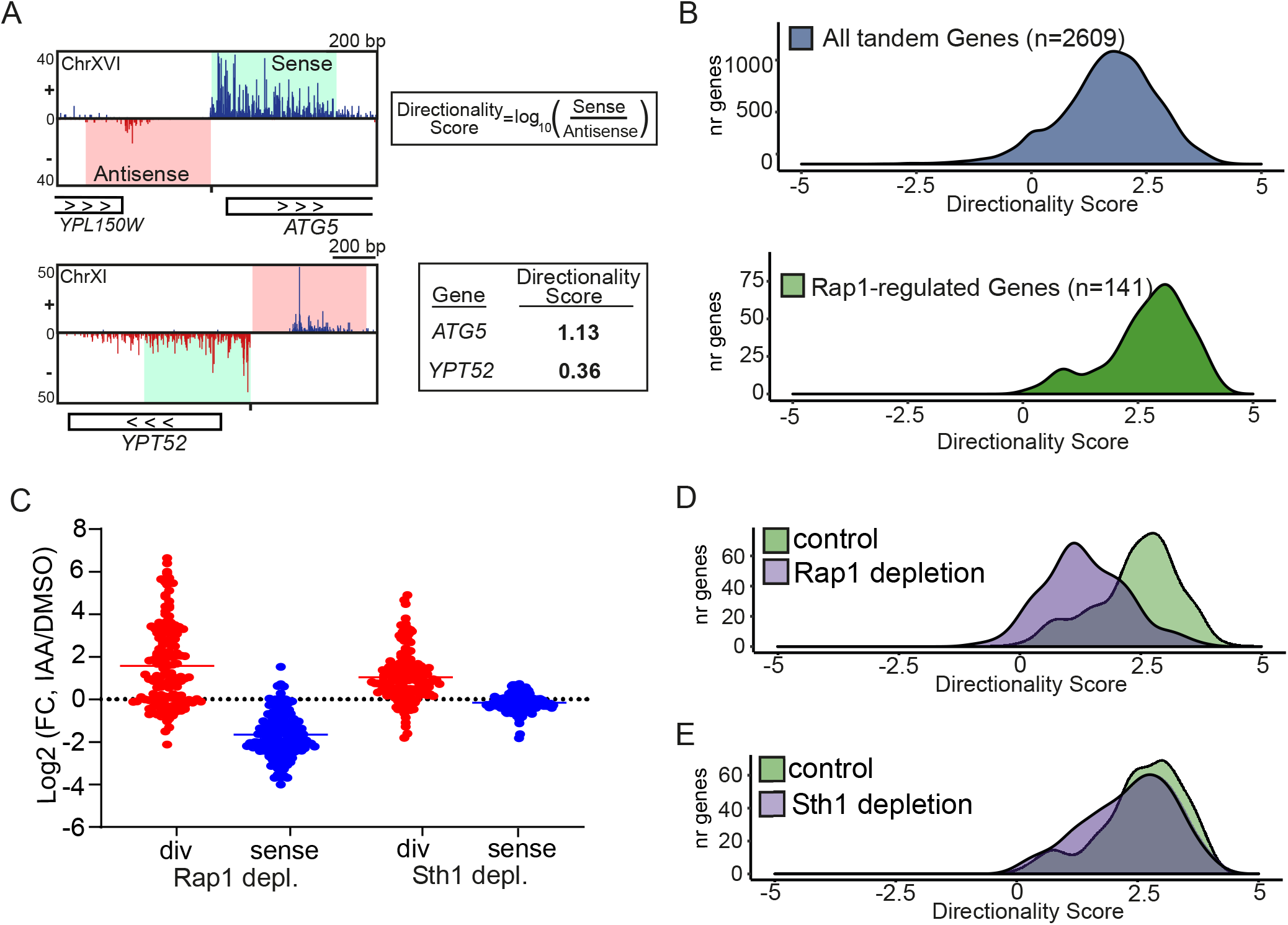
Depletion of RSC increases divergent transcription and alters promoter directionality. **(A)** Approach for calculating directionality score for each promoter. In short, the nascent RNA-seq signals 400 bp upstream (antisense strand) and 400 bp downstream (sense strand) of the coding gene TSS were taken. Subsequently, the ratio of sense over antisense signals (log10) was computed, which was defined as the directionality score. A comparable approach was also described in (Jin et al 2017). **(B)** Density plot of directionality scores for tandem non-overlapping genes (n=2609) and Rap1-regulated genes (n=141). (**C**) Relative changes in transcription levels in cells depleted for Rap1 (*RAP1-AID* + IAA), Sth1 (*STH1-AID* + IAA) or both (*RAP1-AID*/*STH1-AID* + IAA) in comparison to the control treated cells (IAA/DMSO) for 141 Rap1-regulated gene promoters. The nascent RNA-seq signals up to 400 bp upstream (antisense strand, labelled as div) and up to 400 bp downstream (sense strand, labelled as sense) of the TSS were taken. **(D)** Density plots representing promoter directionality of control (*RAP1-AID* +DMSO) and Rap1 depleted cells (*RAP1-AID* + IAA) for Rap1-regulated genes (n=141). **(E)** Similar as D, except comparing control (*STH1-AID* + DMSO) and Sth1 depleted (*STH1-AID* + IAA) cells).

### RSC controls promoter directionality at Rap1 regulated promoters

While our analysis using nascent RNA-seq cannot determine absolute effects in transcription changes, it is possible to dissect relative changes. Promoter directionality score is a suitable relative measurement as it is defined by the ratio between transcription in the sense direction over the divergent direction (Figure 2A) (Jin et al 2017). We computed the directionality score for tandem gene promoters, based on the TSS annotation of a dataset described previously (Park et al 2014). We focused our analysis on Rap1 regulated gene promoters (141 genes) and all tandem, non-overlapping gene promoters where the upstream gene is positioned in the tandem orientation (n=2609 promoters), to avoid confounding signals from divergent coding genes (Figures 2B and S2). As expected, Rap1-regulated genes are highly directional when compared to all tandem genes (Figure 2B). Upon Rap1 depletion, directionality was strongly reduced due to both the increased in the divergent transcription and a decrease in sense transcription (Figures 2C and 2D). Upon Sth1 depletion, the promoter directionality scores were reduced. Indeed, Sth1-depleted cells displayed a relative increase in divergent transcription (Figures 2B and 2D). Thus, at the highly directional Rap1-regulated genes, RSC controls promoter directionality by a repressive effect on divergent transcription.

### RSC controls promoter directionality genome-wide

Divergent noncoding transcription is an inherent feature of gene regulation across species (Jacquier 2009, Pelechano & Steinmetz 2013, Seila et al 2009). Accordingly, many classes of noncoding transcripts have been identified. These include: cryptic unstable transcripts (CUTs) which are processed by the nuclear exosome complex, Xrn1-sensitive antisense non-coding RNA (XUTs), Nrd1-unterminated transcripts (NUTs) and stable unannotated transcripts (SUTs) (Neil et al 2009, Schulz et al 2013, van Dijk et al 2011, Xu et al 2009). These noncoding transcripts (CUTs/SUTs/XUTs/NUTs) are often found at gene promoters and expressed in the divergent direction but are also found at antisense transcription units which overlap protein-coding genes. We examined how expression levels of CUTs/SUTs/XUTs/NUTs were affected upon RSC depletion. While protein-coding gene transcription was relatively decreased (298 genes upregulated vs 567 genes downregulated (fold change (FC)>2, padj<0.05)), noncoding transcription (CUTs/SUTs/XUTs/NUTs) was substantially up-regulated (1330 transcripts upregulated vs 492 transcripts downregulated (FC >2, padj<0.05)) (Figures 3A). Employing the directionality analysis described in Figure 2A, we found that more than 900 displayed a relative increase (FC> 2) in divergent transcription, out of the 2609 tandem and non-overlapping genes (Figure 3B and 3C). Consequently, more than 1000 gene promoters showed a decrease (FC> 2) in directionality score upon Sth1 depletion (S*TH1-AID* + IAA) compared to WT or control cells (*STH1-AID* +DMSO) (Figure 3C). Notably, control cells showed a decreased directionality score compared to the WT, indicating that the AID tag on Sth1 had a small negative impact on RSC function, possibly underestimating the overall effect of RSC (Figure 3C, S3A and S3B). We conclude that RSC is important for limiting divergent noncoding transcription and promoting sense coding transcription at least one third of tandem gene promoters.

**Figure 3.**
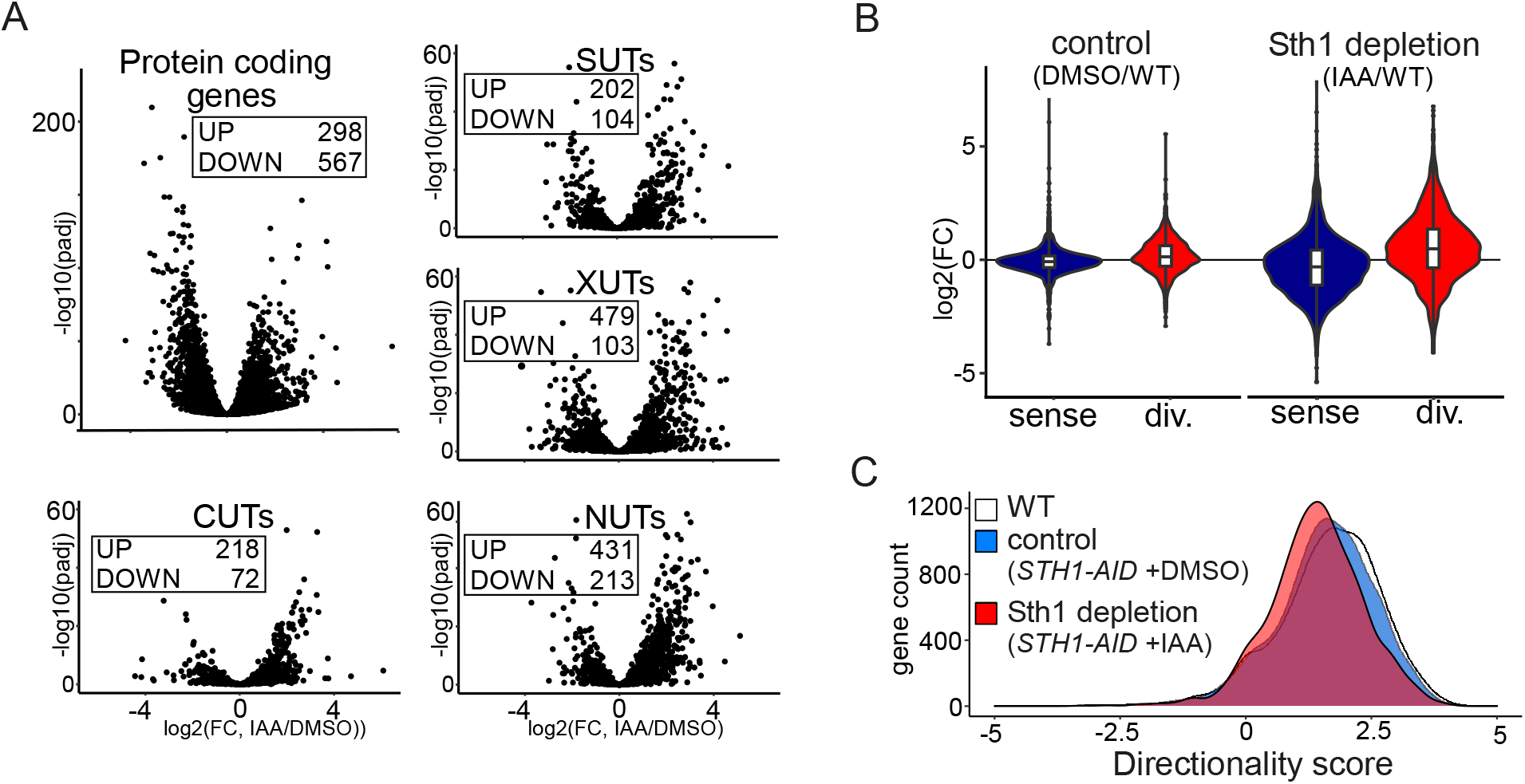
RSC depletion affects promoter directionality genome-wide. **(A)** Volcano plots of nascent-RNA seq data comparing Sth1 depleted to control cells (IAA/DMSO) for different classes of transcription units. Displayed are plots for protein-coding genes, and noncoding transcripts: CUTs, SUTs, XUTs, and NUTs. Number of loci with significantly increased (UP) or decreased (DOWN) levels of transcription are displayed (*padj*<0.05 and log2FC>1). **(B)** Violin plots displaying fold change (log2FC) in nascent-RNA-seq signal, in the sense and divergent direction for control (*STH1-AID* + DMSO) versus WT cells, and Sth1 depletion (*STH1-AID* + IAA) versus WT cells. **(C)** Promoter directionality score of tandem and non-overlapping protein-coding genes in WT, control (*STH1-AID* + DMSO) and Sth1 depleted (*STH1-AID* +IAA) cells.

### RSC-mediated nucleosome organization may limit TBP association upstream of directional promoters

To further explore RSC’s role in controlling divergent transcription, we examined whether there are features that could explain the effect of RSC on promoter directionality. For example, it is known that RSC acts on highly transcribed gene promoters and coding sequences (Biernat et al 2021, Kubik et al 2015, Rawal et al 2018). Our data indicate that the ratio of divergent over sense transcription levels change upon Sth1 depletion, indicating that RSC’s role in promoter directionality is, at least in part, mediated through controlling divergent transcription (Figure 3B and S3A). We divided gene promoters into quintiles (Q1-Q5) based on directionality score in WT cells, wherein Q5 represents the group of promoters with the highest directionality score and Q1 the lowest. Subsequently, we computed the changes in divergent and sense transcription, as well as changes in directionality score for each quintile (Figure 4A). Promoters with the highest directionality score (Q5) in WT cells showed the largest relative increase in divergent transcription after Sth1 depletion (Figure 4A and S4A, left panel), suggesting that RSC-mediated repression of divergent transcription is more prominent at highly directional promoters (Figures 4A and S4A right panel). RSC depletion also affected directionality of gene promoters with the highest sense transcription levels (Q5, sense transcription in WT), albeit less compared to the promoters with highest directionality (Q5, directionality score in WT) (Figure S4B). Control cells (*STH1-AID* +DMSO) displayed a marginal increase in divergent transcription in the most highly expressed and most directional promoters (Q5 sense transcription levels and Q5 directionality in WT, respectively) compared to WT cells, which was consistent with the observation that the AID tag partially affected RSC activity and had a small negative impact on promoter directionality (Figures 3D and S4C). Thus, RSC mainly represses divergent transcription at promoters that are highly directional.

**Figure 4.**
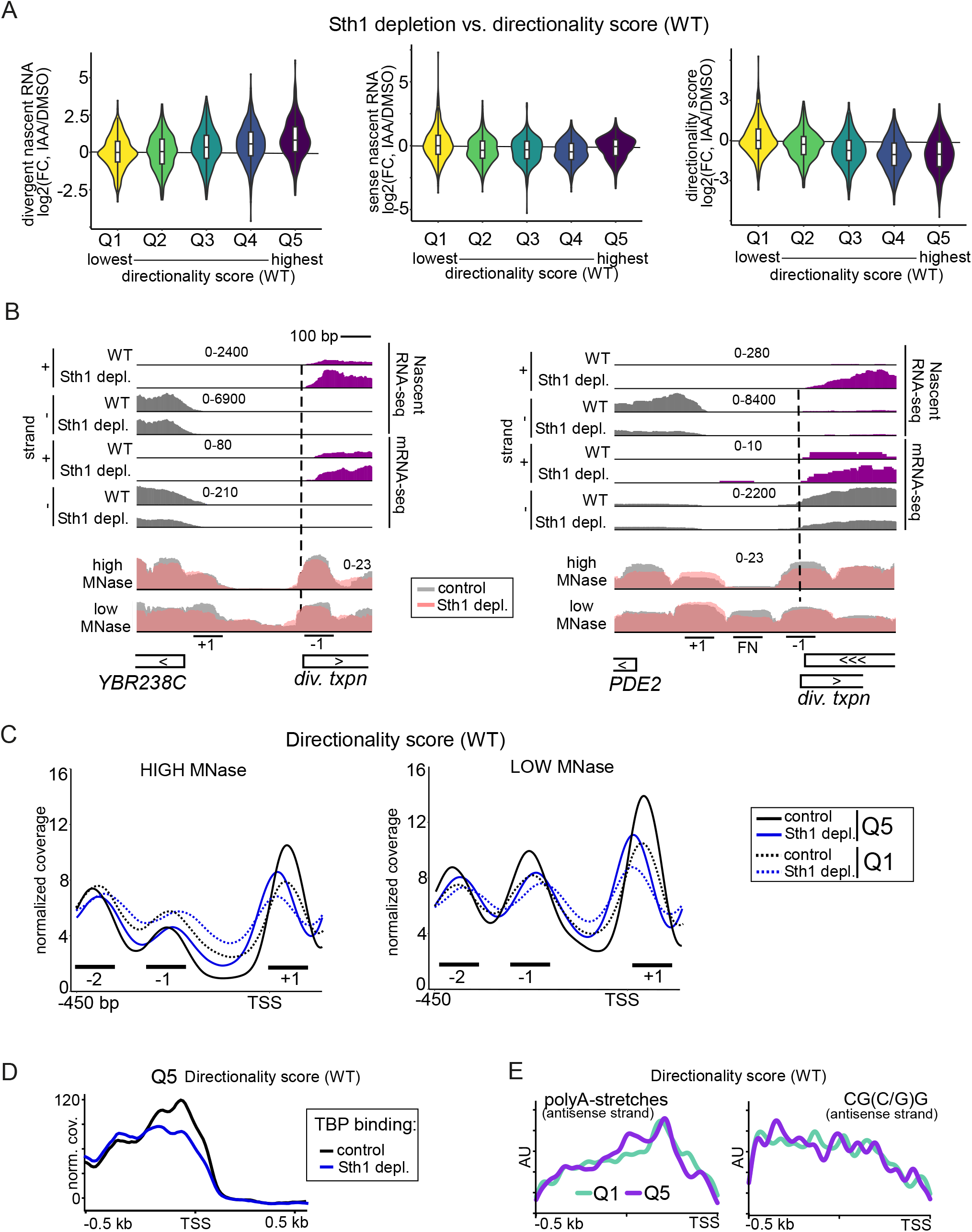
RSC acts on highly directional promoters with fragile nucleosomes. **(A)** Violin plots display log2 of fold change (FC) in divergent transcription (left), sense transcription (centre), and directionality score (right) in Sth1 depleted versus control cells (IAA/DMSO). Tandem genes were sorted on directionality score, and gene groups were divided in five groups (Q1 to Q5). Q1 represents the group of gene promoters with lowest directionality score in WT cells, and Q5 represents the group with the highest directionality score. The range for log10 directionality scores for each group are the following: Q1(−3 −0.9), Q2(0.9 - 1.5), Q3(1.5 - 2), Q4(2 - 2.5), Q5(2.5 - 4.2). **(B)** Example loci that display increased divergent transcription and changes in chromatin structure upon Sth1 depletion. Shown are the data from nascent RNA-seq, mRNA-seq and MNase-seq. **(C)** Metagene gene analysis of MNase-seq data for Q1 and Q5 described in A. Data were centred on coding gene TSS, and displayed are MNase-seq signals for chromatin extracts treated with low MNase or high MNase concentrations. The MNase-seq data were obtained from (Kubik et al 2015, Kubik et al 2018). Marked are the regions representing the +1, −1, and −2 nucleosome positions. **(D)** Metagene analysis of TBP ChIP-seq for Q5 group of gene promoters sorted on directionality score. **(E)** Analysis of RSC complex sequence motifs (A-tracks and CG(C/G)G) in Q1 and Q5 groups of gene promoters sorted on directionality score. Displayed is the distribution in the motif signal on the antisense strand only.

RSC plays a prominent role in positioning nucleosomes in promoters, which in turn affects PIC recruitment and TSS scanning (Hartley & Madhani 2009, Klein-Brill et al 2019). To investigate the mechanisms underlying these effects of RSC on promoter directionality, we assessed the chromatin structure Sth1 depleted cells (Figure 4B and 4C). Specifically, we compared profiles of MNase sensitivity in cells treated with high and low amounts of MNase using a published dataset (Kubik et al 2015). The extraction of nucleosome particles using different MNase concentrations identifies the presence of so-called fragile nucleosomes (Brahma & Henikoff 2019, Kubik et al 2015, Kubik et al 2018). These nucleosome particles appear when extracts are treated with low concentrations of MNase and likely represent partially unwrapped nucleosomes, but have been also suggested to represent chromatin bound non-histone proteins (Brahma & Henikoff 2019, Chereji et al 2017, Kubik et al 2015, Kubik et al 2018, Yan et al 2018).

We observed the RSC-dependent differences in chromatin structure at individual gene promoters. For example, the *PDE2* and *YBR238C* promoters display increased divergent transcription and a small but notable decrease in MNase-seq signal for the −1 nucleosomes in Sth1 depleted cells (Figure 4B). Notably, the *YBR238C* promoter also harbours a fragile nucleosome (Figure 4B, labelled “FN”). The broad effect of Sth1 depletion on nucleosome positioning (−2, −1, and +1) in directional promoters may explain why both divergent and sense transcription are often affected upon RSC depletion.

To determine whether the effect at the *PDE2* and *YBR238C* gene promoters renders genome-wide changes, we analysed nucleosome positioning in the promoter classes sorted by directionality score. First, we found that the group of gene promoters with the highest directionality score (Q5, WT) displayed more defined nucleosome peaks, narrower in width at the +1, −1 and −2 positions compared to gene promoters with the lowest directionality score (Q1) (Figure 4C, left panel). The −1 and −2 nucleosomes were present in the regions where divergent transcription initiates, while +1 nucleosome was present in the region where transcription initiation from the coding TSS occurs. Second, upon depletion of Sth1, nucleosome positioning became broader and less defined for the −1 and −2 positions, while the +1 nucleosome shifted more upstream. Third, the +1, −1 and −2 nucleosomes were more sensitive to MNase concentration in Q5 promoters than Q1 promoters (Figure 4C, right panel), suggesting a prevalence for partially unwrapped nucleosomes in this group. We also compared the effect of promoter directionality on a subset of promoters with comparable sense coding gene transcription levels (Figure S4D and S4E). This analysis revealed that the upstream −1 nucleosome is more defined and sensitive to MNase concentrations for the promoters with the highest directionality score (highest 20%) compared to group of promoters in the lowest 20% directionality score group within this subset of promoters with comparable coding gene transcription, further supporting the idea that positioning of upstream nucleosomes mediates promoter directionality (Figure S4E).

The increase in divergent transcription, as observed in Sth1 depleted cells, is possibly a consequence of altered PIC formation at divergent promoters. We assessed whether association of TATA-binding protein (TBP) was affected upon Sth1 depletion in the group of gene promoters with highest directionality (Q5) using a published TBP ChIP-seq dataset (Kubik et al 2018). Despite that the ChIP-seq data does not have high spatial resolution, we found that upon Sth1 depletion, TBP binding was less affected, perhaps slightly increased, in the region approximately 400 bp upstream of the coding gene TSS which overlaps with the region where core promoters of divergent transcripts are found (Figure 4D). TBP enrichment near the coding TSS decreased, which is expected because RSC promotes TBP recruitment to coding gene promoters (Kubik et al 2015).

Lastly, we examined whether DNA sequence motifs associated with RSC activity are differentially enriched and/or distributed between promoters with low and high directionality. Both polyA stretches and the CG(C/G)G motif are associated with RSC binding (Badis et al 2008, Krietenstein et al 2016, Kubik et al 2018, Lorch et al 2014). We examined the distribution of these sequences in both the sense and antisense strand (Figure 4E and S4F). We found that polyA stretches and the CG(C/G)G motif were enriched in the divergent antisense direction in the group of promoters with the highest directionality (Q5) compared to the group of promoters with lowest directionality (Q1) (Table S1). Interestingly, the A track on the antisense strand showed two peaks in promoters with highest the directionality, possibly indicating that multiple RSC localisations may facilitate the positioning of FNs and −1 nucleosomes at highly directional promoters (Figure 4E). Altogether, these data further support a model where RSC is recruited to distinct positions to promote positioning of upstream nucleosomes in promoters, thereby limiting divergent transcription initiation. The highly positioned nucleosomes, in turn, can form physical barriers which inhibit recruitment of the PIC and RNA Pol II.

### RSC and GRFs are enriched at highly directional promoters

In addition to nucleosome remodeling by RSC, GRFs such as Rap1 are key players for controlling promoter architecture and regulating gene transcription (Brahma & Henikoff 2019, Kubik et al 2017, Kubik et al 2018). To assess how RSC and the GRFs Abf1 and Reb1 associate with respect to promoter directionality, we analysed a published dataset and compared groups of gene promoters with the lowest or highest directionality score (Q1 vs Q5, Figure 4A) (Brahma & Henikoff 2019). In line with the role of RSC in controlling divergent transcription, we found that RSC was more enriched in highly directional gene promoters (compare Q5 to Q1, Figure 5A and S5). Moreover, we found that Abf1 and Reb1 were also more enriched for the group of promoters with the highest directionality score (compare Q5 to Q1, Figure 5A and S5). This suggests that, apart from Rap1, other GRFs may contribute to repressing divergent transcription. To further investigate this hypothesis, we examined a published dataset that measured RNA crosslinked to RNA Pol II (RNA Pol II CRAC) upon Reb1 depletion (Candelli et al 2018b, Zentner et al 2015). We found that depletion of Reb1 increased levels of divergent RNA associated with Pol II notably at the *BAP2* and *CSG2* promoters (Figure 5B).

**Figure 5.**
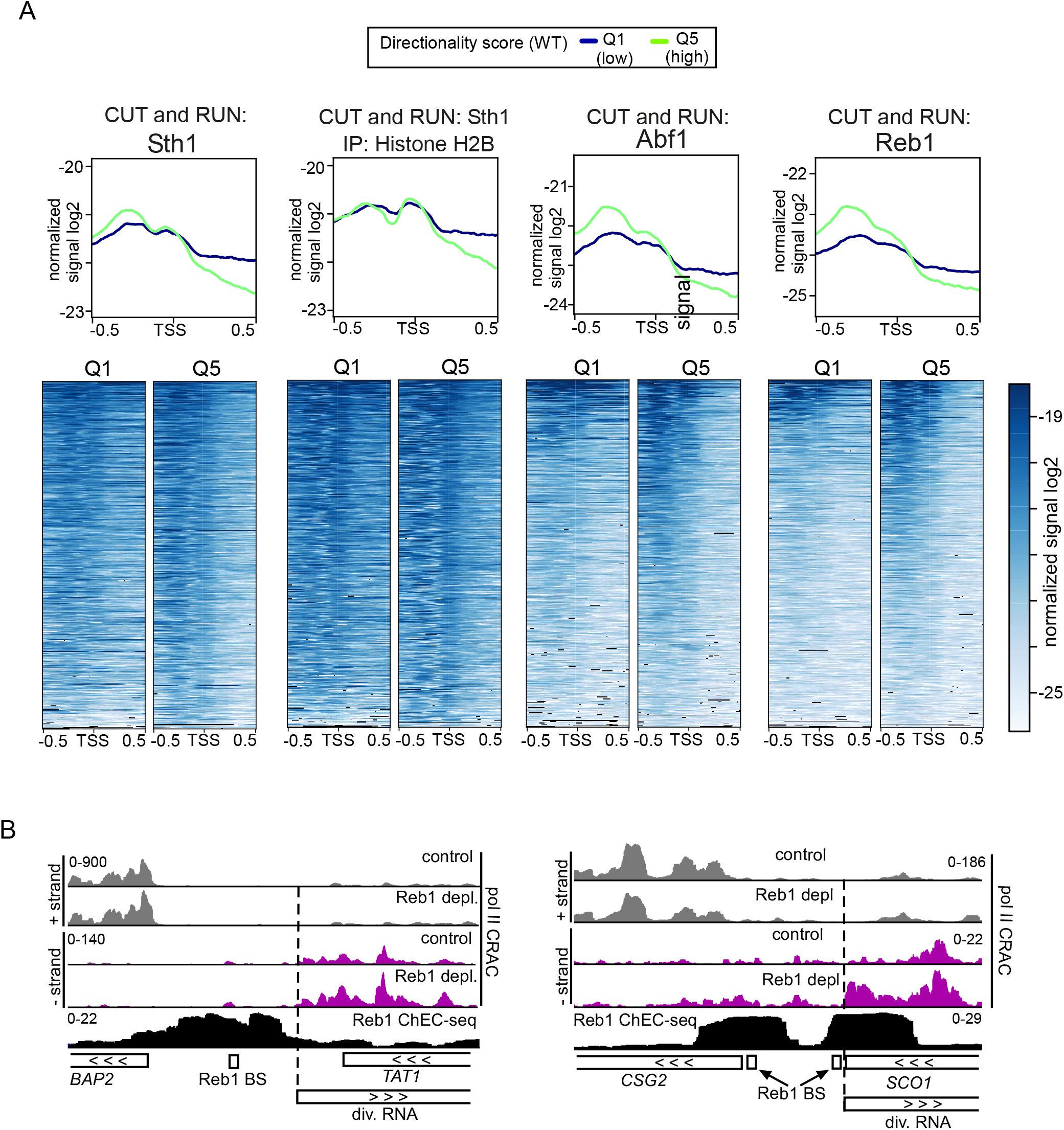
RSC and GRFs are enriched upstream in promoters with a high directional score. **(A)** Metagene analysis and heatmaps of CUT&RUN data of Sth1 (RSC), and the GRFs Abf1 and Reb1. In addition, signals of Sth1 CUT&RUN followed by histone H2B immunoprecipitation are displayed. The data were obtained from (Brahma & Henikoff 2019). A comparison between Q1 (blue), the group of gene promoters with lowest directionality score, and Q5 (green), the group with the highest directionality score, was made. Corresponding heatmaps for Q1 and Q5 are displayed below the graph. The data represents one of n=2 biological repeats. **(B)** Data from RNA Pol II CRAC and Reb1-ChEC-seq datasets described previously (Candelli et al 2018a, Zentner et al 2015), representing RNA Pol II transcription in presence or absence of Reb1 and binding of Reb1, respectively. Displayed are the *BAP2* and *SCO1* loci, which show increased divergent transcription upon Reb1 depletion. Also shown are the positions of the Reb1 binding sites (BS).

### Targeting GRFs or dCas9 to divergent core promoters is sufficient to repress divergent noncoding transcription

Our analysis suggests that Abf1 and Reb1 can repress divergent transcription, possibly using a similar mechanism as Rap1 (Figure 5A). As we previously demonstrated, Rap1 limits initiation of divergent noncoding transcription by occupying target motifs nearby or adjacent to cryptic divergent core promoters (Wu et al 2018). To investigate whether Reb1, Abf1, and transcription factors (TFs) including GRFs can repress divergent transcription, we introduced ectopic transcription factor binding sites adjacent to a divergent core promoter. We used an established fluorescent reporter assay based on the *PPT1* promoter which normally has a noncoding transcript *SUT129* in the divergent direction, with both the coding and the divergent noncoding core promoters driving the transcription of fluorescent reporter genes encoding mCherry and YFP, respectively (Marquardt et al 2014, Wu et al 2018). We cloned the binding site sequences of several transcription factors 20 base pairs (bp) upstream of the *SUT129*/YFP TSS in the *PPT1* reporter (Figure 6A). We selected GRFs (Cbf1, Abf1, and Reb1) and TFs (Gal4, Gcn4, Cat8, Gcr1) that have well-defined DNA binding sequence motifs. To establish that the changes in the reporter signal were dependent on the transcription factor’s presence in the cells and not solely due to the alteration in the underlying promoter DNA sequence, we also measured the reporter signal after deleting or depleting the same transcription factors. For example, for the reporter construct with Reb1 binding sites we measured YFP/mCherry levels in Reb1 control and depleted cells using the auxin induced degron (*REB1-AID* +IAA), whereas for the reporter construct with Cbf1 binding sites we measured the YFP/mCherry levels in WT and *cbf1*Δ cells.

**Figure 6.**
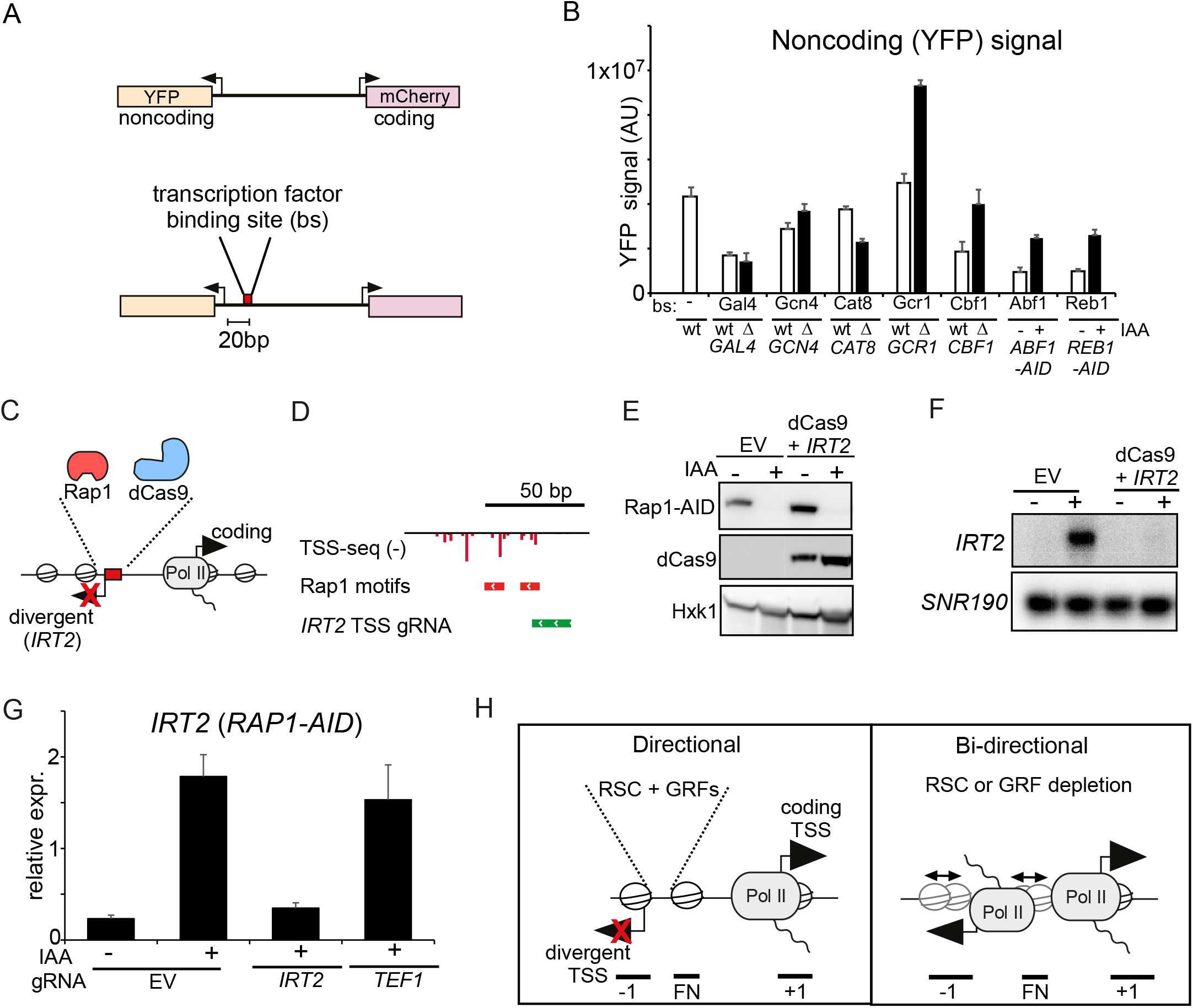
Targeting GRFs or dCas9 to divergent core promoters is sufficient to repress divergent noncoding transcription. **(A)** Schematic overview of the reporter construct. The transcription factor binding sites were cloned 20 nucleotides upstream of the YFP (*SUT129*) TSS. **(B)** YFP, noncoding, signal for constructs harbouring binding sites for Gal4, Gcn4, Cat8, Gcr1, Cbf1, Abf1 and Reb1 cloned into the WT *PPT1* promoter (FW6407). The YFP activity was determined in WT control cells and in gene deletion strains of matching transcription factor binding site reporter constructs (FW6404, FW6306, FW6401, FW6300, FW6402, FW6302, FW6403, FW6424, FW6405, and FW6315). For Abf1 and Reb1 reporter constructs, Abf1 and Reb1 were depleted using the auxin inducible degron (*ABF1*-AID and *REB1*-AID) (FW6415 and FW6411). Cells were treated for 2 hours with IAA or DMSO. Displayed is the mean signal of at least 50 cells. The error bars represent 95% confidence intervals. **(C)** Scheme depicting targeting dCas9 to repress the divergent transcript *IRT2*. **(D)** Design of gRNAs targeting the *IRT2* TSS. Displayed are TSS-seq data from after Rap1 depletion (Wu et al 2018). **(E)** Western blot of Rap1 and dCas9 detected with anti-V5 and anti-FLAG antibodies, respectively. Cells harbouring *RAP1*-AID were treated with IAA to deplete Rap1 and induce *IRT2* expression in either cells with empty vector (EV) or cells harbouring a dCas9 construct and a construct expressing the gRNA targeted to the Rap1 binding site (FW8477, FW8531). Hxk1 was used as a loading control, detected using anti-Hxk1 antibodies. (**F**) *IRT2* expression detected by northern blot for strains and treatments described in E. As a loading control, the membrane was re-probed for *SNR190.* **(G)** *IRT2* expression as detected in by RT-qPCR of as described in D. A control gRNA was included in the analysis targeted to the TSS of *TEF1* (FW8527). Displayed are the mean signals of n=3 biological repeats and SEM. **(H)** Model for promoter directionality.

Our data indicate that several GRFs were able to repress transcription in the divergent direction and increase promoter directionality, suggesting that they behave similarly to Rap1 (Figure 6B) (Wu et al 2018). Specifically, *cbf1*Δ cells and depletion of Abf1 and Reb1 (*ABF1-AID* or *REB1-AID* +IAA) resulted a reduced signal in divergent noncoding transcription compared to control cells (Figure 6B). Introduction of binding site motifs for one transcription factor, Gcr1, did not affect noncoding (YFP) direction relative to the WT, but the YFP signal increased in *gcr1*Δ cells suggesting that Gcr1 also can repress divergent transcription (Figure 6A and 6B). The TFs that did not negatively affect the divergent noncoding transcript signal (Gal4, Gcn4, and Cat8) were possibly less active under the growth conditions where we performed the reporter assay (Figure 6B). For example, Gal4 and Gcn4 are most active under galactose containing medium and under amino acid starvation, respectively. Notably, Gal4 and Gcn4 showed increased signal in the coding direction, suggesting that they are still somewhat active in the conditions tested but perhaps not sufficiently active for repressing divergent transcription (Figure S6A). We conclude that, as we reported previously for Rap1 (Wu et al 2018), other transcription factors can repress divergent transcription when targeted to regions near a divergent gene promoter.

Our data indicate that RSC-mediated nucleosome positioning and GRFs can repress initiation of divergent transcription, and thereby promote directionality of gene promoters (Challal et al 2018, Wu et al 2018). One explanation is that these proteins are barriers for PIC formation and Pol II recruitment. If true, then any protein stably associated with DNA near divergent TSSs should be able to interfere with divergent transcription. To test this, we targeted an exogenous protein to a divergent TSS. We used the catalytically inactivated version of *S. pyogenes* Cas9 (dCas9), which is a bulky protein (over 150 kDa) and has been widely used as tool to modulate transcription and chromatin (Gilbert et al 2013). We induced divergent transcription by depleting Rap1 and used gRNAs to target dCas9 to the core promoter of *IRT2*, which is divergently transcribed from *RPL43B* and strongly induced upon Rap1 depletion, Figures 6C-E). As expected, Rap1 depletion induced expression of *IRT2* (Figure 6D and 6E). Importantly, we did not detect *IRT2* expression when dCas9 was targeted to the *IRT2* TSS (Figure 6F, 6G and S6B). Expression of *IRT2* was still detectable in cells where dCas9 was targeted to the *TEF1* the promoter or upstream in the *MLP1* promoter (*iMLP1*) (Figure 6G and S6B). We conclude that steric hindrance by DNA-binding proteins (such as dCas9) is sufficient for repression of divergent transcription.

## Discussion

Transcription from promoters is bidirectional, leading to sense coding and divergent noncoding transcription. However, the directionality (sense over divergent transcription) of promoters greatly varies indicating that there are mechanisms in place to control directionality of promoters (Jin et al 2017). Why some promoters are more directional than others is not well understood. Here, we showed that RSC mediated nucleosome positioning represses divergent noncoding transcription. Furthermore, we provided evidence that a wide range of DNA-binding proteins, including nucleosomes, GRFs, and dCas9, are capable of repressing divergent transcription. We propose that such RSC-positioned nucleosomes and other DNA-binding factors form physical barriers at promoters which prevent transcription in the divergent direction, thereby promoting promoter directionality.

### RSC-mediated nucleosome positioning represses divergent transcription

The RSC complex largely contributes to generating NDRs by ‘pushing’ the nucleosomes positioned at +1 and −1 positions ‘outwards’, widening the NDR (Ganguli et al 2014, Hartley & Madhani 2009, Parnell et al 2008). RSC also binds directly to DNA via A-track sequence motifs (Krietenstein et al 2016, Kubik et al 2018, Lorch et al 2014). How does RSC mediate the position of the −1 nucleosome to repress divergent transcription? Interestingly, we found that A-tracks were abundant in promoters on both strands indicating that directional positioning of nucleosomes upstream in promoters may contribute to preventing divergent transcription. RSC also promotes the partial unwinding of nucleosomes, which has been also referred to as “fragile” nucleosomes as these nucleosome particles are sensitive to MNase concentration (Brahma & Henikoff 2019, Kubik et al 2018). These sub-nucleosome complexes are also bound by GRFs. We found that at directional promoters the −1 nucleosome and FN are more defined compared to promoters with less directionality, suggesting that the formation of sub-nucleosome complexes controls promoter directionality (Figure 6H). We further found that depletion of RSC resulted in less defined consensus nucleosome patterns and reduced sensitivity to MNase. We propose that nucleosomes strongly positioned by RSC inhibit PIC formation and Pol II recruitment for the divergent direction (Figure 6C). Upon RSC depletion, the positioning of the −1 nucleosome and FN is less strong and PIC formation can occur.

RSC-mediated nucleosome positioning also functions in suppressing other aberrant transcription events. For example, RSC represses antisense transcription from the 3’ ends of gene bodies, which is spatially distinct from the divergent noncoding transcription events which we report here (Alcid & Tsukiyama 2014, Cucinotta et al 2021). In addition, RSC-mediated positioning of the +1 nucleosome can affect TSS selection, resulting in increased transcription initiation in the sense direction within promoters (Cucinotta et al 2021, Klein-Brill et al 2019, Kubik et al 2019). This work demonstrates that RSC is important for repressing divergent noncoding transcription upstream in coding gene promoters, and this is important for controlling the directionality of promoters. Thus, RSC has a widespread function in promoting coding gene transcription and limiting aberrant noncoding transcription in each of the divergent, antisense, and sense directions.

The opening of chromatin controlled by RSC facilitates the recruitment of sequence-specific TFs to promoters (Floer et al 2010). It may be possible that GRFs or other non-histone proteins mediate the repression of divergent transcription via RSC, as has been proposed by others (Brahma & Henikoff 2019, Chereji et al 2017). Our analysis identified examples of divergent transcription events in the same promoter which were repressed by Rap1 and RSC, but utilised distinct TSSs, suggesting that RSC can repress divergent transcription independently of GRFs (Figures 1D and 6H). How other GRFs act with RSC in controlling divergent transcription remains to be dissected further. Based on our analysis, it appears more likely that RSC and GRFs act together to repress divergent transcription and promote transcription in the protein-coding direction.

### Model for promoter directionality

Nucleosomes form barriers for transcription. In the context of repressing divergent transcription, chromatin assembly factor I (CAF-I) plays a widespread role in yeast, demonstrating that chromatin assembly upstream in promoters is important in repressing divergent transcription (Marquardt et al 2014). On the other hand, opening of chromatin in promoters increases divergent transcription. For example, increased histone lysine 56 acetylation (H3K56ac) leads to more divergent transcription in yeast (Marquardt et al 2014). Conversely, deacetylation of histone H3 by Hda1 deacetylase complex (Hda1C) facilitates repression of divergent transcription (Gowthaman et al 2021). The role of RSC at promoters is likely different from CAF-I, H3K56ac, and Hda1C. We propose that RSC-mediated positioning of NDR-flanking nucleosomes in promoters generates barriers for limiting RNA Pol II recruitment and initiation of divergent transcription. In line with this, differential TSS usage has been observed in RSC-depleted cells (Kubik et al 2019).

In a previous work we showed that Rap1 can repress divergent transcription at gene promoters likely by occupying cryptic divergent promoters (Wu et al 2018). Our new analysis suggests that other GRFs have potentially the same ability in a model system. Moreover, we found that Abf1 and Reb1 were more enriched at highly directional promoters, suggesting that these GRFs also perform this role *in vivo*. dCas9 has the same capacity to repress aberrant noncoding transcription when targeted to divergent core promoters. We propose that proteins physically interfere with divergent transcription when bound to DNA. With this view, RSC activity is important to position crucial nucleosomes and to ensure that cryptic divergent promoters are ‘protected’ and transcriptionally repressed. Thus, GRFs and positioned nucleosomes constitute essential components for promoter organization, which promote sense coding transcription and limit divergent noncoding transcription (Figure 6H). While our studies focussed on tandem gene promoters. It is worth nothing that sequence specific DNA binding factors, such as Tbf1 and Mcm1, have the ability insulate two independently regulated divergent gene promoter pairs from each other, thus effectively repress divergent transcription (Yan et al 2015). Additionally, in mammalian cells the multifunctional transcription factor CTCF directly represses initiation divergent noncoding transcription at hundreds for promoters, indicating that the mechanism of DNA binding factors repressing divergent transcription is likely conserved and widespread (Luan et al 2021).

RSC is part of the SWI/SNF family of chromatin remodellers related to the mammalian PBAF and BAF complexes (Mohrmann & Verrijzer 2005). The BAF complex in mouse embryonic stem cells (esBAF), has related functions in repressing noncoding transcription to some extent (Hainer et al 2015). Like RSC, esBAF positions the nucleosomes flanking the NDR. Depletion of esBAF leads to less stable nucleosome positioning, and consequently to widespread increased noncoding transcription, of which some are divergent noncoding transcription events. We anticipate that the role of RSC and BAF complexes in controlling divergent transcription and promoter directionality is likely conserved.

Further work will be required to dissect the interplay between GRFs and chromatin states in controlling aberrant noncoding transcription and promoter directionality. These regulatory complexes are present and conserved across eukaryotes. Thus, our study may provide important insights on how promoter directionality is controlled in multi-cellular organisms.

## Materials and Methods

### Strains and plasmids and growth conditions

Strains isogenic to the Saccharomyces cerevisiae BY4741 strain background (derived from S288C) were used for this investigation. A one-step tagging procedure was used to generate endogenous carboxy (C)-terminally tagged auxin-inducible degron (AID) alleles of Rap1 and Sth1 (Nishimura et al 2009). The *RAP1-AID* strain was previously described, which contains three copies of the V5 epitope and the IAA7 degron (Wu et al 2018). For tagging *RPB3* with the FLAG epitope, we used a one-step tagging procedure using a plasmid (3xFLAG-pTEF-NATMX-tADH1, gift from Jesper Svejstrup). The single copy integration plasmids for expression of *Oryza sativa TIR1* (*osTIR1*) ubiquitin E3 ligase were described previously (Wu et al 2018).

For auxin-inducible degron (*AID*) depletion experiments, induction of AID-tagged protein depletion was performed with 3-indole-acetic acid (IAA) (Sigma-Aldrich) during the exponential growth phase (approx. OD_600_ 0.8). The *pPPT1-pSUT129* (*pPS*, mCherry-YFP) fluorescent reporter plasmid was described previously (gift from Sebastian Marquardt, University of Copenhagen) (Marquardt et al 2014). The target motifs for each transcription factor (obtained from the YeTFaSCo database (de Boer & Hughes 2012)) were introduced by blunt-end cloning into unique restriction site (SspI) proximal to the *SUT129* TSS and verified by Sanger sequencing. After linearisation by digestion with EcoRI, the reporter construct was transformed and integrated to replace the endogenous *PPT1-SUT129* locus.

For CRISPR interference (CRISPRi) experiments, yeast expression constructs for nuclease-inactivated Cas9 (D10A/H840A mutations) from *Streptococcus pyogenes* –dCas9 (Addgene #46920) (Gilbert et al 2013, Qi et al 2013) – were sub-cloned into single copy integration plasmids by restriction cloning. The 3xFLAG epitope was introduced by Gibson-style cloning (NEBuilder HiFi, NEB) at the C-terminus of the construct, in-frame with the ORF. dCas9-3xFLAG expression construct on single copy integration plasmids was transformed into yeast after linearisation with PmeI. Single guide RNA (sgRNA) plasmids (gift from Elçin Ünal, UC Berkeley) (Chen et al 2017) were generated by site-directed mutagenesis (Q5 Site-Directed Mutagenesis Kit, NEB), using specific primers containing the sgRNA 20-mer target sequence (split in half across the two primers). sgRNA target sequence selection was guided by availability of “NGG” protospacer adjacent motif (PAM) sites and transcription start-site sequencing (TSS-seq) data (Wu et al 2018). sgRNA expression plasmids were integrated at the *SNR52* locus after linearisation by XbaI digestion. Strains, plasmids, and oligonucleotide sequences are listed in Table S2, S3, and S4, respectively.

### RNA extraction

Yeast cells were harvested from cultures for RNA extraction by centrifugation, then washed once with sterile water prior to snap-freezing in liquid nitrogen. RNA was extracted from frozen yeast cell pellets using Acid Phenol:Chloroform:Isoamyl alcohol (125:24:1, Ambion) and Tris-EDTA-SDS (TES) buffer (0.01 M Tris-HCl pH 7.5, 0.01 M EDTA, 0.5% w/v SDS), by rapid agitation (1400 RPM, 65 °C for 45 minutes). After centrifugation, the aqueous phase was collected and RNA was precipitated at −20 °C overnight in ethanol with 0.3 M sodium acetate. After centrifugation and washing with 80% (v/v) ethanol solution, dried RNA pellets were resuspended in DEPC-treated sterile water and subsequently stored at −80 °C.

### RT-qPCR

Reverse transcription and quantitative PCR for *IRT2* performed as described previously (Tam & van Werven 2020) (Moretto et al 2018). In short, total RNA was purified using the NucleoSpin RNA kit (Macherey-Nagel) according to the manufacturer’s instructions. For reverse transcription, ProtoScript II First Strand cDNA Synthesis Kit (New England BioLabs) was used and 500 ng of total RNA was provided as template in each reaction. qPCR reactions were prepared using EXPRESS SYBR GreenER SuperMix (Thermo Fisher Scientific) and *IRT2* levels were quantified on Applied Biosystems 7500 Fast Real-Time PCR System (Thermo Fisher Scientific). The result of n=3 biological replicates are shown. Signals were normalised over *ACT1*. Primer sequences are listed in Table S4.

### Western blot

Western blots were performed as described previously (Chia et al 2017, Chiu et al 2018). Protein extracts were prepared from whole cells after fixation with trichloroacetic acid (TCA). Samples were pelleted by centrifugation and incubated with 5% w/v TCA solution at 4 °C for at least 10 minutes. Samples were washed with acetone, pelleted, and dried. Samples were then resuspended in protein breakage buffer (50 mM Tris (pH 7.5), 1 mM EDTA, 2.75 mM dithiothreitol (DTT)) and subjected to disruption using a Mini Beadbeater (Biospec) and 0.5 mm glass beads. Protein extract samples were mixed with SDS-PAGE sample buffer (187.5 mM Tris (pH 6.8), 6.0% v/v β-mercaptoethanol, 30% v/v glycerol, 9.0% v/v SDS, 0.05% w/v Bromophenol blue) in a 2:1 ratio by volume, and protein samples were denatured at 95 °C for 5 minutes.

SDS-PAGE (polyacrylamide gel electrophoresis) was performed using 4-20% gradient gels (Bio-Rad TGX) and samples were then transferred onto PVDF membranes by electrophoresis (wet transfer in cold transfer buffer: 3.35% w/v Tris, 14.9% w/v glycine, 20% v/v methanol). Membranes were incubated in blocking buffer (1% w/v BSA, 1% w/v non-fat powdered milk in phosphate buffered saline with 0.01% v/v Tween-20 (PBST) buffer) before primary antibodies were added to blocking buffer for overnight incubation at 4 °C. For probing with secondary antibodies, membranes were washed in PBST and anti-mouse or anti-rabbit IgG horseradish peroxidase (HRP)-linked antibodies were used for incubation in blocking buffer (1 hour, room temperature). Signals corresponding to protein levels were detected using Amersham ECL Prime detection reagent and an Amersham Imager 600 instrument (GE Healthcare).

### Antibodies

The following antibodies were used for western blotting.

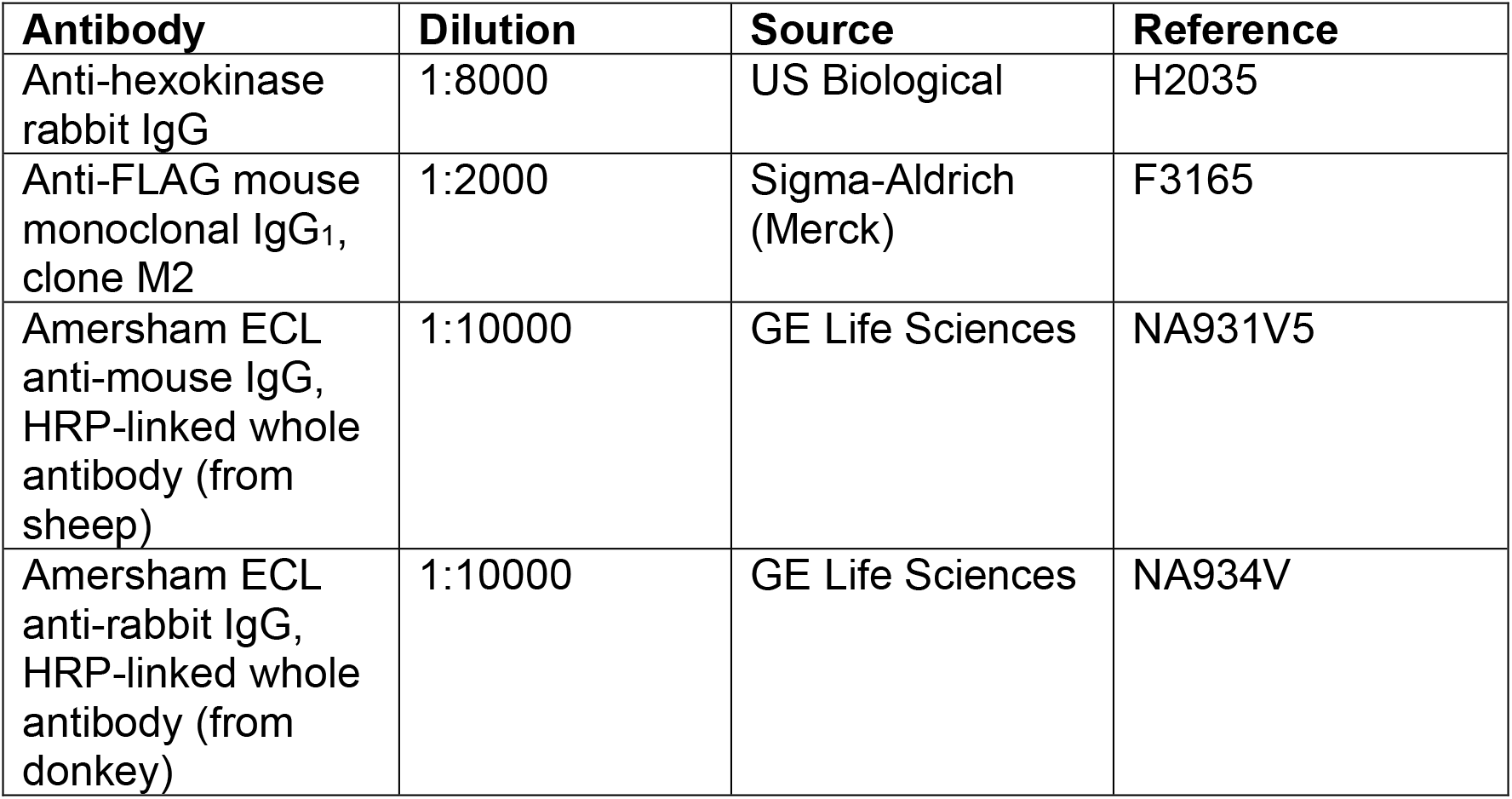

| Antibody | Dilution | Source | Reference |
| --- | --- | --- | --- |
| Anti-hexokinase rabbit IgG | 1:8000 | US Biological | H2035 |
| Anti-FLAG mouse monoclonal IgG<sub>1</sub>, clone M2 | 1:2000 | Sigma-Aldrich (Merck) | F3165 |
| Amersham ECL anti-mouse IgG, HRP-linked whole antibody (from sheep) | 1:10000 | GE Life Sciences | NA931V5 |
| Amersham ECL anti-rabbit IgG, HRP-linked whole antibody (from donkey) | 1:10000 | GE Life Sciences | NA934V |

### Northern blot

Northern blots were performed as described previously (Chia et al 2017, Wu et al 2018). Briefly, RNA samples (10 μg per lane) were incubated in sample denaturation buffer (1 M deionised glyoxal, 50% v/v DMSO, 10 mM sodium phosphate (NaP_i_) buffer (pH 6.8)) at 70 °C for 10 minutes, loading buffer (10% v/v glycerol, 2 mM NaP_i_ buffer, 0.4% w/v bromophenol blue) was added, and RNA samples were subjected to electrophoresis (2 hours at 80 V) on an agarose gel (1.1% v/v agarose, 0.01 M NaP_i_ buffer). Capillary transfer was used to transfer total RNA onto positively charged nylon membranes (GE Amersham Hybond N+). Bands corresponding to mature rRNA were visualised by staining with methylene blue solution (0.02% w/v methylene blue, 0.3 M sodium acetate).

The nylon membranes were incubated for at least 3 hours at 42 °C in hybridisation buffer (1% w/v SDS, 40% v/v deionised formamide, 25% w/v dextran sulfate, 58 g/L NaCl, 200 mg/L sonicated salmon sperm DNA (Agilent), 2 g/L BSA, 2 g/L polyvinyl-pyrolidone, 2 g/L Ficoll 400, 1.7 g/L pyrophosphate, 50 mM Tris pH 7.5) or ULTRAhyb Ultrasensitive Hybridization Buffer (Thermo Fisher Scientific) to minimise non-specific probe hybridisation. Probes were synthesised using a Prime-it II Random Primer Labeling Kit (Agilent), 25 ng of target-specific DNA template, and radioactively labelled with dATP [α-32P] (Perkin-Elmer or Hartmann Analytic). The oligonucleotides used to amplify target-specific DNA templates for *IRT2* and *SNR190* northern blot probes by PCR can be found in Table S4.

Blots were hybridised overnight with radioactively labelled probes at 42 °C, and then washed at 65 °C for 30 minutes each with the following buffers: 2X saline-sodium citrate (SSC) buffer, 2X SSC with 1% w/v SDS, 1X SSC with 1% SDS, and 0.5X SSC with 1% SDS. For image acquisition, membranes were exposed to storage phosphor screens before scanning on the Typhoon FLA 9500 system (GE Healthcare Life Sciences). To strip membranes prior to re-probing for different transcripts, membranes were washed with stripping buffer (1 mM Tris, 0.1 mM EDTA, 0.5% w/v SDS) at 85 °C until negligible residual signal remained on the blots.

### Fluorescence microscopy

Yeast cells were grown in YPD media (small batch cultures) to the exponential growth phase and fixed with formaldehyde (3.7% w/v), incubating at room temperature for 15 minutes. Fixed cells were washed with phosphate-sorbitol buffer (0.1 M KP_i_ (pH 7), 0.05 M MgCl_2_, 1.2 M sorbitol), and resuspended in the same buffer prior to imaging. Images were acquired using a Nikon Eclipse Ti-E inverted microscope imaging system (Nikon) equipped with a 100x oil objective (NA 1.4), SOLA SE light engine (Lumencor), ORCA-FLASH 4.0 camera (Hamamatsu) and NIS-Elements AR software (Nikon). 500 ms exposure time was specified and GFP and mCherry filters were used to detect YFP and mCherry signals, respectively.

To quantify whole cell fluorescence signals, measurements were performed using ImageJ (version 1.52i, NIH) (Schneider et al 2012) for the YFP and mCherry channels. Regions of interest (ROIs) were manually drawn around the border of each cell. Mean signal represents the mean intensity in each channel per cell multiplied by the cell area, and the signal for YFP and mCherry was corrected for cell-free background fluorescence in a similar manner. WT cells without integrated fluorescent reporter plasmids were also measured to determine auto-fluorescence signal. 50 cells were quantified for each sample.

### RNA-seq with *S. pombe* spike in

Total RNA was extracted from *S. cerevisi*ae pellets spiked in with *S. pombe* cells in a 10:1 ratio of *S. cerevisiae*:*S. pombe* as described above, treated with rDNase in solution (Machery-Nagel), and purified by spin column (Machery-Nagel) prior to preparation of sequencing libraries. 500 ng of RNA input material was used for polyadenylated (polyA) RNA sequencing. Libraries were prepared using TruSeq Stranded mRNA kit (Illumina) according to the manufacturer’s instructions (10 or 13 PCR cycles). The libraries were multiplexed and sequenced on either the HiSeq 2500 or 4000 platform (Illumina), and generated ∼45 million 101 bp strand-specific paired-end reads per sample on average.

### Nascent RNA sequencing (Nascent RNA-seq)

For nascent RNA sequencing (nascent RNA-seq), RNA fragments associated with RNA polymerase II subunit Rpb3 endogenously tagged with 3xFLAG epitope at the C-terminus were isolated by affinity purification as described previously (Churchman & Weissman 2011; 2012, Moretto et al 2021). Small batch cultures of yeast cells grown in YPD media were collected by centrifugation, the supernatant was aspirated, and cell pellets were immediately snap-frozen by submerging in liquid nitrogen to minimise changes in nascent transcription activity (e.g. in response to cell resuspension in cold lysis buffer with high concentrations of salts and detergents). Frozen cell pellets were dislodged from centrifuge tubes and stored at −80 °C. Cells were subjected to cryogenic lysis by freezer mill grinding under liquid nitrogen (SPEX 6875D Freezer/Mill, standard program: 15 cps for 6 cycles of 2 minutes grinding and 2 minutes cooling each). Yeast “grindate” powder was stored at −80 °C. 2 g of yeast grindate was resuspended in 10 mL of 1X cold lysis buffer (20 mM HEPES pH 7.4, 110 mM potassium acetate, 0.5% v/v Triton-X-100, 1% v/v Tween-20) supplemented with 10 mM MnCl_2_, 1X Roche cOmplete EDTA-free protease inhibitor, and 50 U/mL SUPERase.In RNase Inhibitor. Chromatin-bound proteins were solubilised by incubation with 1320 U of DNase I (RQ1 RNase-free DNase I, Promega) on ice for 20 minutes. Lysates containing solubilised chromatin proteins were clarified by centrifugation at 20,000 x *g* for 10 minutes (at 4 °C), and the supernatant was taken as input for immunoprecipitation using 500 μL of anti-FLAG M2 affinity gel suspension (A2220, Sigma-Aldrich) per sample (2.5 hours at 4 °C). After immunoprecipitation, the supernatant was removed and beads were washed 4 times with 10 mL of cold wash buffer each time (1X lysis buffer with 50 U/mL Superase.In RNase inhibitor and 1 mM EDTA). After the last wash, agarose beads were transferred to small chromatography spin columns (Pierce Spin Columns, Thermo Fisher), and competitive elution of protein complexes containing Rpb3-3xFLAG protein from the resin was performed by incubating beads with 300 μL of elution buffer (1X cold lysis buffer with 2 mg/mL 3xFLAG peptide) for 30 minutes at 4 °C (3xFLAG peptide provided by the Peptide Chemistry Science Technology Platform, The Francis Crick Institute). Elution was performed twice and 600 μL of eluate was subjected to acid phenol:chloform RNA extraction and ethanol precipitation. A significant amount of 3xFLAG peptide co-precipitates with the RNA as a contaminant and is later removed by spin column purification. Purified RNA was fragmented to a mode length of ∼200 nucleotides using zinc ion-mediated fragmentation (Ambion AM870, 70 °C for 4 minutes). Fragmented RNA was purified using miRNeasy spin columns (miRNeasy mini kit, QIAGEN), which retain RNAs approximately 18 nucleotides or more in length. Purified RNA was quantified by Qubit (Thermo Fisher) and approximately 150 ng of RNA was subjected to rRNA depletion using the Ribo-Zero Gold rRNA Removal Kit (Yeast) (Illumina MRZY1324, now discontinued). Libraries were prepared using the TruSeq Stranded Total RNA kit (Illumina) according to the manufacturer’s instructions (14 PCR cycles). The libraries were multiplexed and sequenced on the HiSeq 4000 platform (Illumina), and generated ∼28 million 101 bp strand-specific paired-end reads per sample on average.

### RNA-seq data analysis

Adapter trimming was performed with cutadapt (version 1.9.1) (Martin 2011) with parameters “--minimum-length=25 --quality-cutoff=20 -a AGATCGGAAGAGC -A AGATCGGAAGAGC”. BWA (version 0.5.9-r16) (Liao et al 2014) using default parameters was used to perform the read mapping independently to both the *S. cerevisiae* (assembly R64-1-1, release 90) and *S. pombe* (assembly ASM294v2, release 44) genomes. Genomic alignments were filtered to only include those that were primary, properly paired, uniquely mapped, not soft-clipped, maximum insert size of 2kb and fewer than 3 mismatches using BamTools (version 2.4.0; (Barnett et al 2011)). Read counts relative to protein-coding genes were obtained using the featureCounts tool from the Subread package (version 1.5.1) (Liao et al 2014). The parameters used were “-O --minOverlap 1 --nonSplitOnly --primary -s 2 -p -B -P -d 0 -D 1000 -C --donotsort”.

Differential expression analysis was performed with the DESeq2 package (version 1.12.3) within the R programming environment (version 3.3.1) (Love et al 2014). The spiked-in *S. pombe* transcripts were used to assess the size factors, potentially mitigating any impact a global shift in the *S. cerevisiae* read counts would have on DESeq2’s usual normalisation procedure.

### Nascent RNA-seq data analysis

Adapter trimming was performed with cutadapt (version 1.9.1) (Martin 2011)) with parameters “--minimum-length=25 --quality-cutoff=20 -a AGATCGGAAGAGC -A AGATCGGAAGAGC -u -50 -U “50”. The RSEM package (version 1.3.0) (Li & Dewey 2011) in conjunction with the STAR alignment algorithm (version 2.5.2a) (Dobin et al 2013) was used for the mapping and subsequent gene-level counting of the sequenced reads with respect to all *S. cerevisiae* genes downloaded from the Ensembl genome browser (assembly R64-1-1, release 90; (Kersey et al 2016)). The parameters used were “--star-output-genome-bam --forward-prob 0”, and all other parameters were kept as default. Differential expression analysis was performed with the DESeq2 package (version 1.12.3) (Love et al 2014) within the R programming environment (version 3.3.1). An adjusted p-value of <= 0.01 was used as the significance threshold for the identification of differentially expressed genes.

STAR genomic alignments were filtered to include only unspliced, primary, uniquely mapped, and properly paired alignments with a maximum insert size of 500 bp. The featureCounts tool from the Subread package (version 1.5.1) (Liao et al 2014) was used to obtain insert fragment counts within defined intervals (e.g. +100 bp) around the 141 Rap1 binding sites (separate intervals for Watson and Crick strand alignments, 242 intervals total) using the parameters “-O --minOverlap 1 -- nonSplitOnly --primary -s 2 -p -B -P -d 0 -D 600 -C”. Strand-specific reads were only counted if they overlapped with the interval on the corresponding strand. DESeq2 was used to perform differential expression analysis for intervals around Rap1 binding sites as described above, and the DESeq2 size factors calculated with respect to the transcriptome or S. pombe transcriptome were used for normalisation of the per-sample counts. Genomic annotations of CUTs, SUTs, XUTs, and NUTs (various noncoding RNA species) was obtained from (Wery et al 2016) (http://vm-gb.curi.e.fr/mw2/), and filtered for transcripts greater than 200 nt in length to match the insert size distribution of RNA-seq libraries.

### MNase-seq/TBP ChIP-seq/CUT&RUN analysis

Publicly available datasets were obtained from NCBI Gene Expression Omnibus (GEO) MNase-seq (GSE73337, GSE98260) (Challal et al 2018, Kubik et al 2015). Adapters were trimmed from MNase-seq reads using cutadapt as described above. Adapter-trimmed reads were mapped to the *S. cerevisiae* genome (Ensembl assembly R64-1-1, release 90) (Zerbino et al 2018) with BWA (version 0.5.9-r16) (Li & Durbin 2009) using default parameters. Only uniquely mapped and properly paired alignments that had no more than two mismatches in either read of the pair and an insert size 120 – 200 bp were kept for paired-end MNase-seq alignments. To generate genome-wide coverage tracks for nucleosome occupancy from MNase-seq data, the DANPOS2 dpos command (version 2.2.2) (Chen et al 2013) was used with parameters “--span 1 --smooth_width 20 --width 40 --count 1000000”.

TBP ChIP-seq data were obtained from GSE98260 (Kubik et al 2018). CUT&RUN data were obtained from GSE116853 (Brahma & Henikoff 2019). The binding of TBP from the above-mentioned ChIP-seq data and the binding of Sth1/Abf1/Reb1 from the above-mentioned CUT&RUN-ChIP data were plotted for genes belonging to the specified categories with deepTools (version 3.3.0) (Ramirez et al 2016), using the computeMatrix parameters “reference-point --referencePoint TSS --upstream 500 -- downstream 500”. The bedgraph files of CUT&RUN-ChIP data were previously clipped to the size of sacCer3 chromosomes, values were normalised over the sum of each sample and transformed to log2.

### Promoter directionality score analysis

To calculate directionality scores for coding gene promoters, a curated list of coding gene TSSs was obtained from published TSS sequencing data as described (Park et al 2014). Any missing TSS coordinates for coding genes were supplemented with the TSS annotation from Ensembl (assembly R64-1-1, release 90) (Zerbino et al 2018), generating a list of 6,646 *S. cerevisiae* coding gene TSSs. To avoid quantification of divergent direction transcription that constituted coding transcription fo r an upstream divergent gene, overlapping and divergent gene pairs were removed from the analysis resulting in 2,609 non-overlapping and tandem genes. To simplify the counting analysis, the coverage for each paired-end read from nascent RNA-seq was reduced to the single 3’ terminal nucleotide of the strand-specific read using genomeCoverageBed function within BEDTools (version 2.26.0) (Quinlan & Hall 2010) with parameters “-ibam stdin -bg -5 -scale %s -strand %s”. “Sense” direction windows encompassed nucleotide positions +1 to +500 in the coding direction relative to the TSS, and “antisense” direction windows encompassed nucleotide positions -1 to -500 in the divergent direction relative to the TSS. The total number of reads with 3’ end positions falling within the sense and antisense direction windows was quantified for each gene using the computeMatrix tool within deepTools (version 2.5.3) (Ramirez et al 2016) using parameters “reference-point --referencePoint center --upstream 500 --downstream 500 --binSize 1 --scale 1”. A value of 1 was added to all counts to avoid dividing by zero. For each window, the mean read count was calculated from three biological replicate experiments and used for subsequent calculation and plotting.

### Stratification of promoters and sequence analysis

Gene promoters were stratified in quintiles (Q1 to Q5) using Matt (version 1.3.0) (Gohr & Irimia 2019), according to sense transcription levels or directionality score (log10) in the WT strain. Analyses were repeated on only promoters with intermediate gene transcription levels (belonging to Q3 according to sense transcription level). Analysis for enrichment and distribution of A-tracks (AAAAAAA) and GC-rich motifs (CG(C/G)G) as defined in (Kubik et al 2015) was performed with Matt (version 1.3.0), using the *test_regexp_enrich* and *get_regexp_prof* functions (Gohr & Irimia 2019).

### Data plotting and visualisation

Bar plots, scatter plots, and volcano plots were generated using GraphPad Prism (version 7 or 8). Screenshots of sequencing data were captured using the Integrative Genomics Viewer (IGV, Broad Institute, version 2.4.15) (Robinson et al 2011). The RStudio integrated development environment (version 1.0.143) was used within the R statistical computing environment (version 3.4.0) for data analysis and visualisation. Software packages within the tidyverse (version 1.2.1) collection were used for data analysis and plotting. The following functions within ggplot (version 3.0.0) were used for plotting: violin plots, geom_violin with parameters “scale = count”;, geom_boxplot with parameters “outlier.colour = NA”; smoothed density plots, geom_density with default parameters; scatter plots, geom_point with default parameters; marginal density histogram plots, ggMarginal with parameters “type = “histogram”, bins = 40, size = 8). The Cairo graphics library (version 1.17.2) was used to generate heat map plots for the RNA-seq data after fold-change values were calculated for bins of 5 nt within defined intervals, comparing between two samples.

### Statistical analysis

Information regarding any statistical tests used, number of samples, or number of biological replicate experiments is stated in the corresponding figure legends. For Students’ *t*-tests, calculated *p* values less than 0.05 were considered significant. Plotted error bars in individual figures are stated in the figure legend, as either standard error of the mean (SEM) or 95% confidence intervals (CI).

### Publicly available datasets used in this study

MNase-seq and TBP-ChIP seq data were obtained from GSE73337 (Kubik et al 2015) and GSE98260 (Kubik et al 2018), respectively. TSS annotation was described in GSE49026 (Park et al 2014). Rap1 regulated gene promoters were described in (Wu et al 2018). The annotations of CUTs, SUTs, NUTs, XUTs were described in http://vm-gb.curi.e.fr/mw2/ (Neil et al 2009, Schulz et al 2013, van Dijk et al 2011, Xu et al 2009). CUT&RUN data were obtained GSE116853 (Brahma & Henikoff 2019). Pol II CRAC data were described in GSE97913 and GSE97915 (Candelli et al 2018b). Reb1-ChEC-seq data were described in GSE67453 (Zentner et al 2015).

## Author Contributions

F.J.v.W. and A.C.K.W. conceived the project. A.C.K.W. performed most experiments. C.V., H.P. and A.C.K.W. performed the bioinformatic analyses. D.S. and A.C.K.W performed the *S. pombe* normalized mRNA-seq experiment. F.M. performed northern blot in Figure S6B. A.C.K.W. and F.J.v.W. designed the experiments. F.J.v.W. wrote the manuscript with input from the other authors. F.J.v.W. supervised the project.

## Acknowledgements

We thank the Crick Advanced Sequencing and Genomics Equipment Park Facilities for experimental support, and Sebastian Marquardt and Jesper Svejstrup for sharing reagents. This research was funded in whole, or in part, by the Wellcome Trust (FC001203). For the purpose of Open Access, the author has applied a CC BY public copyright licence to any Author Accepted Manuscript version arising from this submission. This work was supported by the Francis Crick Institute (FC001203), which receives its core funding from Cancer Research UK (FC001203), the UK Medical Research Council (FC001203), and the Wellcome Trust (FC001203).

## Data Availability

The accession number for the RNA sequencing data reported in this paper is GSE179256.

## Data Files

**Table S1 to S4**

**Gel scans,** contains the western and northern blot scans used for assembling the figures in this study.

---

**Figure S1.**
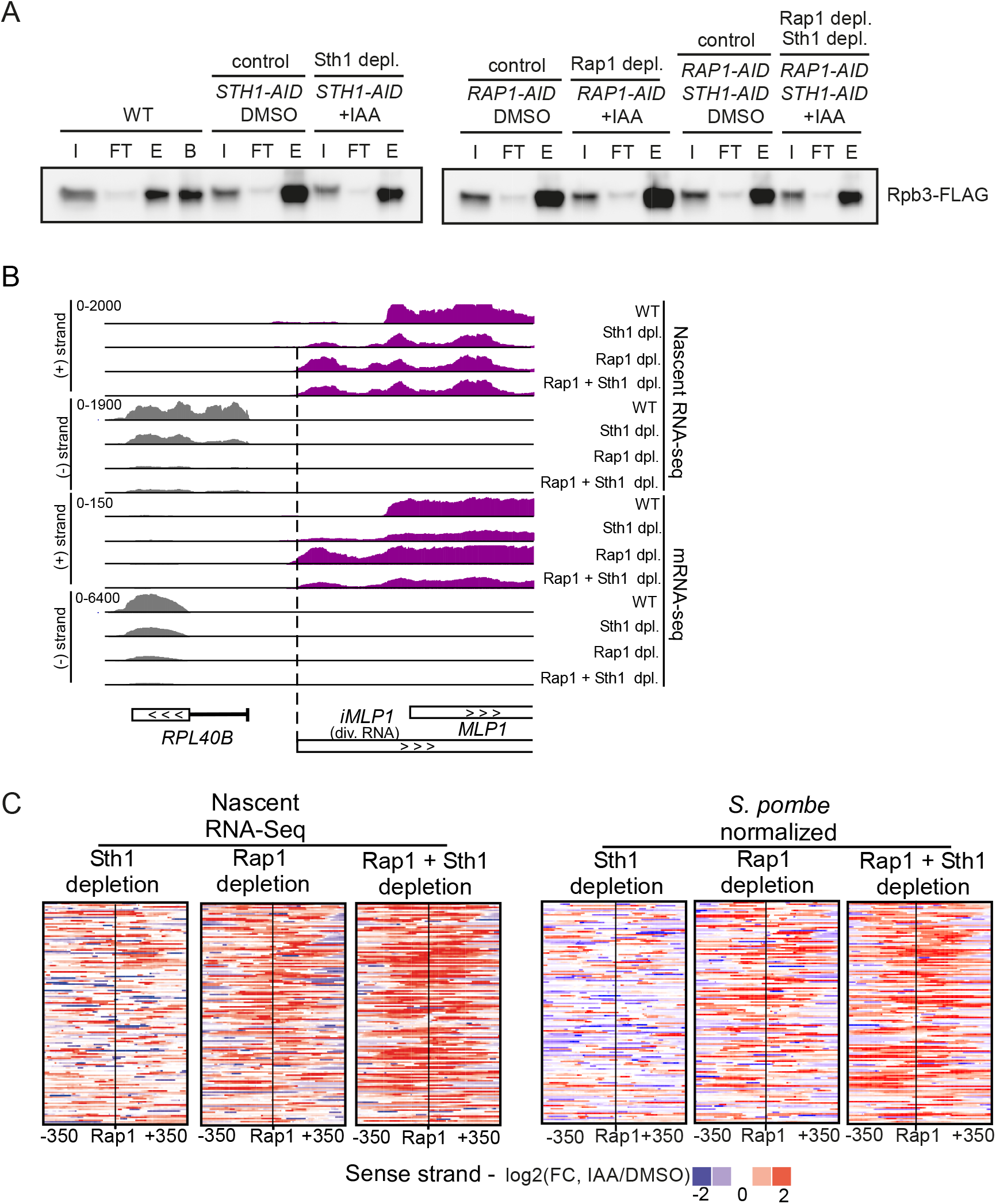
Depletion of RSC increases divergent transcript accumulation at Rap1-regulated genes. **(A)** Rpb3-FLAG immunoprecipitation as detected by western blot. Rpb3-FLAG was immunoprecipitated from wild-type (WT) (FW7228) cells or cells harbouring AID tagged alleles for Sth1, Rap1, and Sth1 + Rap1 (*STH1-AID*, *RAP1-AID*, *RAP1-AID* + *STH1-AID*) (FW7238, FW7220, and FW7232). AID tagged cells were either treated with IAA or DMSO. Samples corresponding to input (I), flow-through (FT), FLAG elution (E), and boiling in sample buffer from FLAG beads (B) are shown. **(B)** Nascent RNA-seq and mRNA-seq signals for *RPL40B* (coding sense direction, grey) and *iMLP1* (divergent coding direction, purple). (**C**) Matching the analysis of Figure 1E, on the sense strand data is displayed. Indicated are the three clusters (c1, c2, and c3) based on previous analysis (Wu et al 2018) for 141 Rap1-regulated gene promoters.

**Figure S2.**
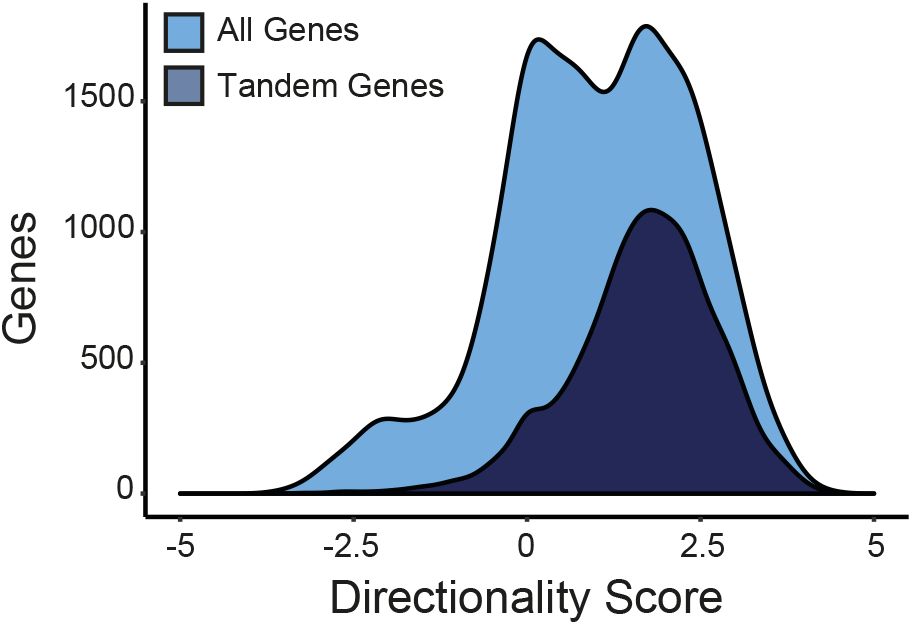
Density plots of promoter directionality scores. Density plots representing promoter directionality scores in WT cells for all genes and tandem genes (n=2609).

**Figure S3.**
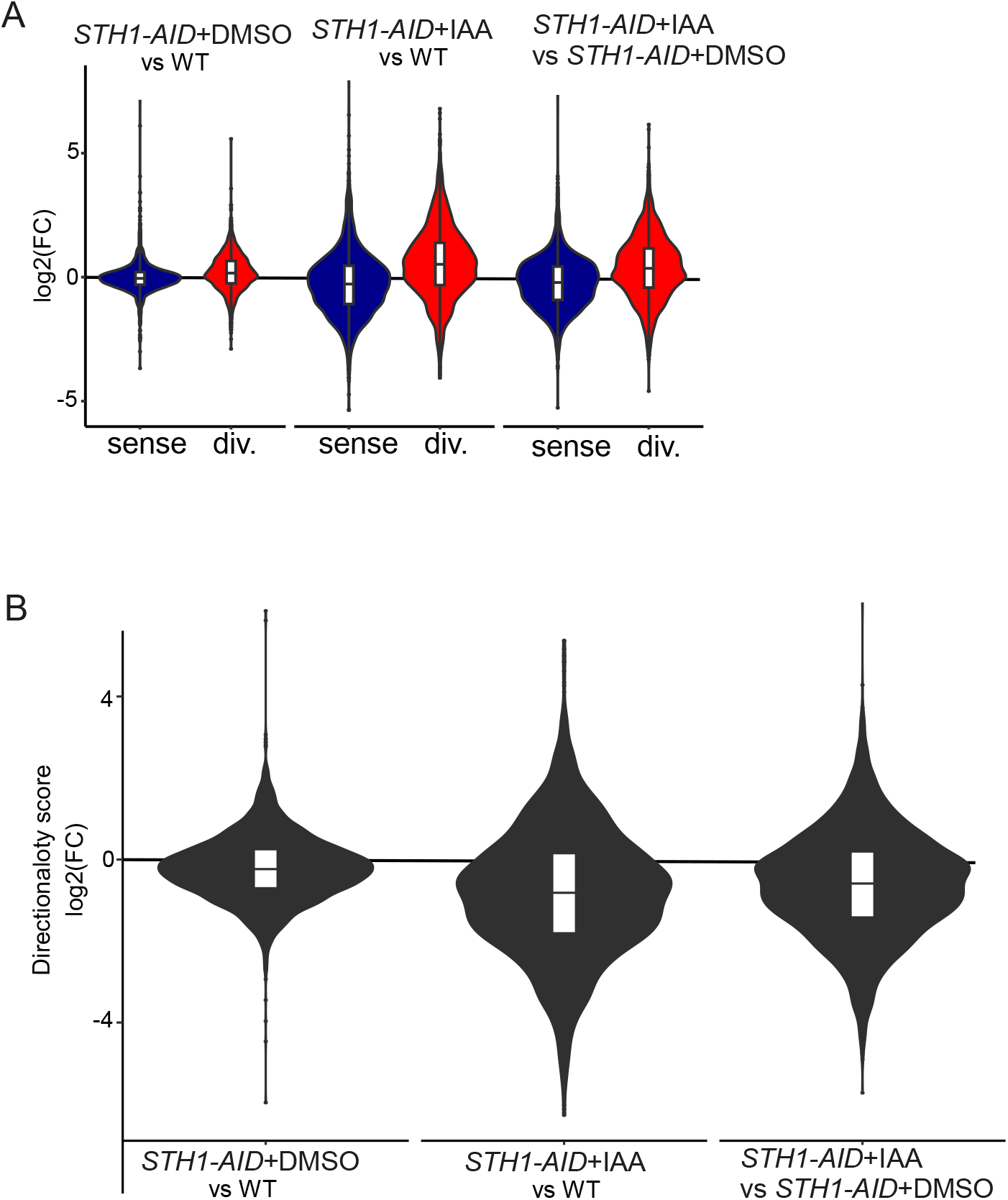
RSC depletion affects promoter directionality genome-wide. **(A)** Violin plots depicting sense and divergent transcription level changes for control (*STH1-AID* + DMSO) over WT, Sth1 depletion (*STH1-AID* +IAA) over WT, and Sth1 (*STH1-AID* +IAA)/ (*STH1-AID* + DMSO) samples. **(B)** Same comparisons as A, but the changes in promoter directionality score are displayed in violin plots.

**Figure S4.**
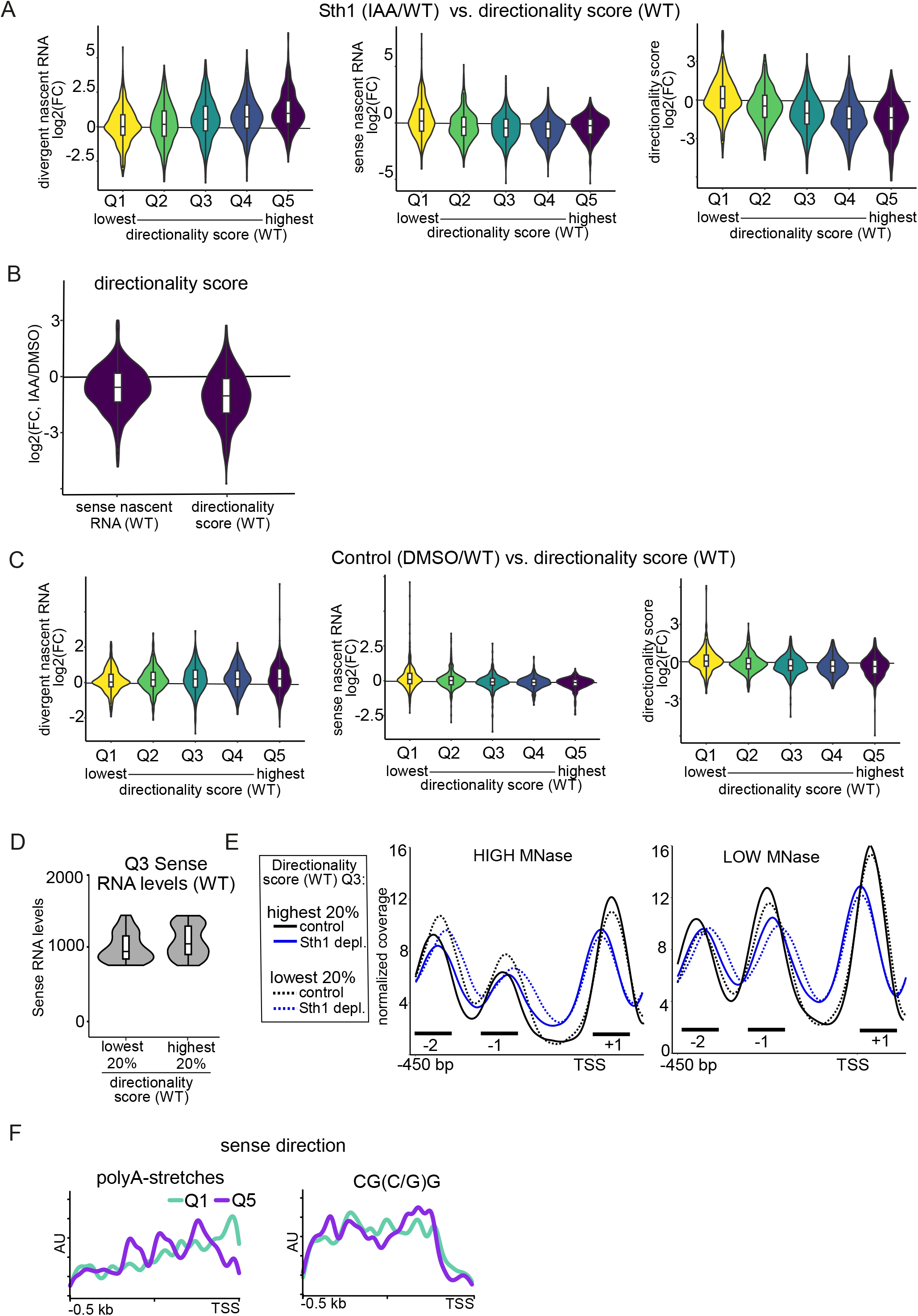
RSC acts at highly directional promoters. **(A)** Violin plots displaying fold change (log2(FC)) in divergent transcription, sense transcription, and directionality score. Tandem genes were sorted on directionality score, and gene groups were divided in five groups (Q1 to Q5). Compared is Sth1 depletion (IAA/WT) sorted on directionality score. The range for log10 directionality scores for each group are the following: Q1(−3 - 0.9), Q2(0.9 - 1.5), Q3(1.5 - 2), Q4(2 - 2.5), Q5(2.5 - 4.2) **(B)** Violin plots displaying log2 fold change (FC) in directionality score of Sth1 depleted versus control (IAA/DMSO) for the group gene promoters (Q5) with highest sense transcription levels or highest directionality score (Q5) in WT cells. **(C)** Similar analysis as A, displayed is the control (DMSO/WT) sorted on directionality score. **(D)** Violin plots of sense direction nascent RNA signals for a group of gene promoters with low and high directionality scores (lowest 20% vs highest 20%), but that have comparable intermediate transcription levels in the sense direction (Q3). Displayed are the sense nascent RNA-seq signals for the genes with 20% lowest directionality score and top 20% directionality score within this select group of gene promoters (n=105). **(E)** Metagene analysis of MNase-seq data of the two groups of gene promoters described D in control and Sth1 depleted cells. **(F)** Distribution of RSC associated DNA sequences in promoters sorted on directionality score. Displayed are the analysis of A-tracks and CG(C/G)G motifs. Distribution of the signals with respect to the sense direction is shown.

**Figure S5.**
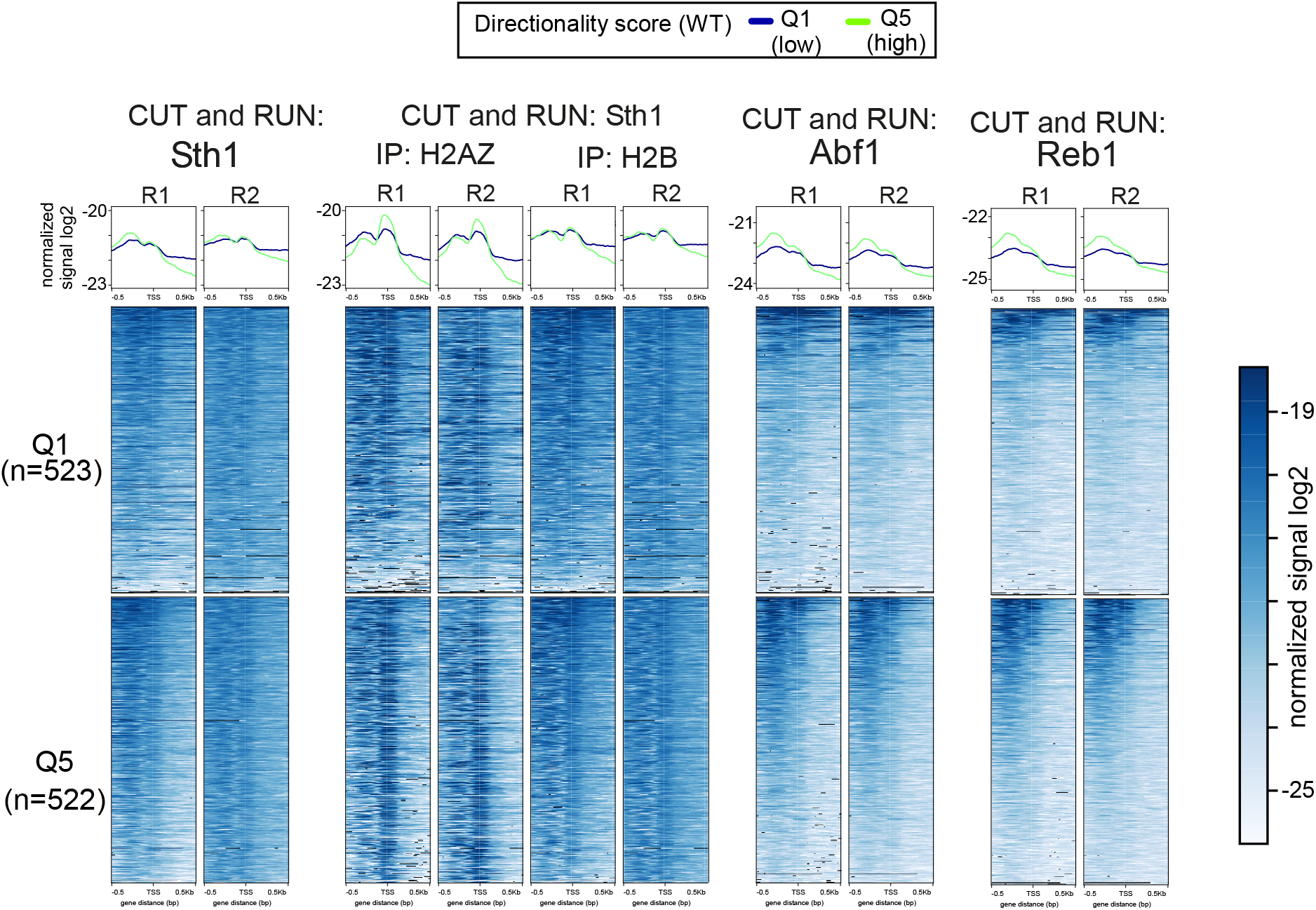
RSC and GRFs are enriched upstream in promoters with a high directionality score. Same analysis as in Figure 5A, except that the two biological repeats are displayed. Metagene analysis and heatmaps of CUT&RUN data of Sth1 (RSC), and the GRFs Abf1 and Reb1 (top). In addition, signals of Sth1 CUT&RUN followed by histone H2B immunoprecipitation are displayed. The data were obtained from (Brahma & Henikoff 2019). Compared Q1 (blue) being the group of gene promoter with lowest directionality score, and Q5 (green) being the group with the highest directionality score. Corresponding heatmaps for Q1 and Q5 are displayed (bottom).

**Figure S6.**
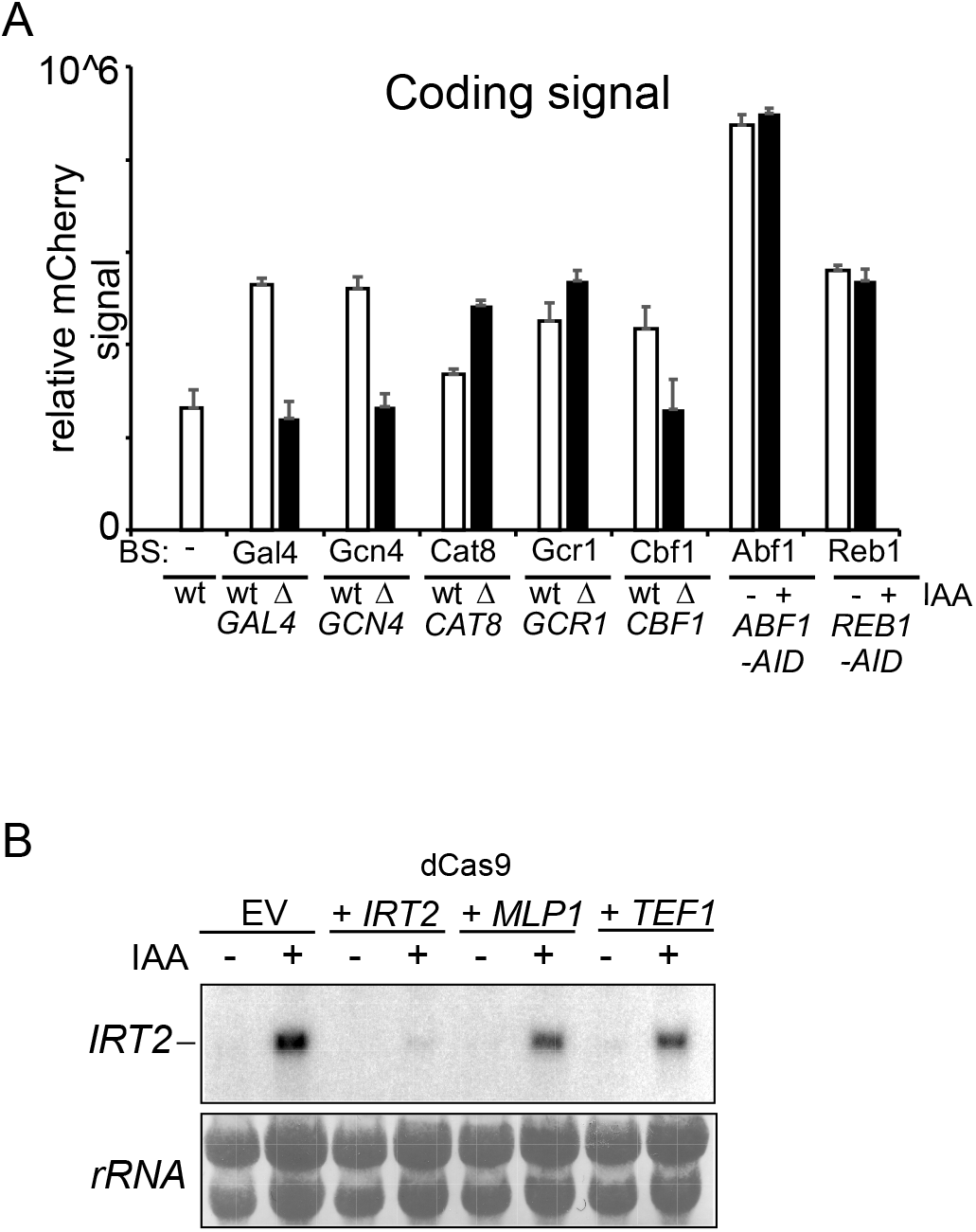
Targeting GRFs or dCas9 to divergent core promoters is sufficient to repress divergent noncoding transcription. **(A)** mCherry (coding direction) signals for constructs harbouring binding sites for Gal4, Gcn4, Cat8, Gcr1, Cbf1, Abf1 and Reb1 cloned to the WT *PPT1* promoter (FW6407). The mCherry activity was determined in WT control cells and in gene deletion stains of matching transcription factor binding site reporter constructs (FW6404, FVW6306, FW6401, FW6300, FW6402, FW6302, FW6403, FW6424, FW6405, and FW6315). For Abf1 and Reb1 reporter constructs, Abf1 and Reb1 were depleted using the auxin inducible degron system (*ABF1*-AID and *REB1*-AID) (FW6415 and FW6411). Cells were treated for 2 hours with IAA or DMSO. Displayed are the mean signal of at least 50 cells. The error bars represent 95% confidence intervals. **(B)** Northern blot of *IRT2* expression in the Rap1 depletion of samples strains described below. Cells harbouring *RAP1-AID* were treated IAA to deplete Rap1 and induce *IRT2* and expressing dCas9 were used for the analysis. Empty vector (EV), and gRNAs targeting *IRT2*, including control gRNAs targeting *iMLP1* and *TEF1* were used (FW8477, FW8531, FW8529, and FW8535). As a loading control rRNA staining is shown.

