## Supplementary material for "RSC and GRFs confer promoter directionality by limiting divergent noncoding transcription": Table S2

**Table S3. Genotypes of strains used in this study**

| Strain Number | Genotype |
|---|---|
| FW7228 | *MATa his3Δ1 leu2Δ0 met15Δ0 ura3Δ0 rpb3::RPB3-3xFLAG::NATMX* |
| FW7220 | *MATa his3Δ1 leu2Δ0 met15Δ0 ura3Δ0 rpb3::RPB3-3xFLAG::NATMX leu2::pGPD1-osTIR1::LEU2 sth1::STH1-V5-AID::KANMX6* |
| FW7232 | *MATa his3Δ1 leu2Δ0 met15Δ0 ura3Δ0 rpb3::RPB3-3xFLAG::NATMX leu2::pGPD1-osTIR1::LEU2 rap1::RAP1-V5-AID::KANMX6 sth1::STH1-V5-AID::KANMX6* |
| FW7238 | *MATa his3Δ1 leu2Δ0 met15Δ0 ura3Δ0 rpb3::RPB3-3xFLAG::NATMX leu2::pGPD1-osTIR1::LEU2 rap1::RAP1-V5-AID::KANMX6* |
| FW6404 | *MATa his3Δ1 leu2Δ0 met15Δ0 ura3Δ0 ppt1::p615-YFP-Cbf1_bs-pPPT1-mCherry::NATMX6* |
| FW6306 | *MATa his3Δ1 leu2Δ0 met15Δ0 ura3Δ0 cbf1::KANMX ppt1::p615-YFP-Cbf1_bs-pPPT1-mCherry::NATMX6* |
| FW6401 | *MATa his3Δ1 leu2Δ0 met15Δ0 ura3Δ0 ppt1::p612-YFP-Gcn4_bs-pPPT1-mCherry::NATMX6* |
| FW6300 | *MATa his3Δ1 leu2Δ0 met15Δ0 ura3Δ0 gcn4::KANMX ppt1::p612-YFP-Gcn4_bs-pPPT1-mCherry::NATMX6* |
| FW6402 | *MATa his3Δ1 leu2Δ0 met15Δ0 ura3Δ0 ppt1::p613-YFP-Cat8_bs-pPPT1-mCherry::NATMX6* |
| FW6302 | *MATa his3Δ1 leu2Δ0 met15Δ0 ura3Δ0 cat8::KANMX ppt1::p613-YFP-Cat8_bs-pPPT1-mCherry::NATMX6* |
| FW6403 | *MATa his3Δ1 leu2Δ0 met15Δ0 ura3Δ0 ppt1::p614-YFP-Gal4_bs-pPPT1-mCherry::NATMX6* |
| FW6424 | *MATa his3Δ1 leu2Δ0 met15Δ0 ura3Δ0 gal4::KANMX ppt1::p614-YFP-Gal4_bs-pPPT1-mCherry::NATMX6* |
| FW6405 | *MATa his3Δ1 leu2Δ0 met15Δ0 ura3Δ0 ppt1::p616-YFP-Gcr1_bs-pPPT1-mCherry::NATMX6* |
| FW6315 | *MATa his3Δ1 leu2Δ0 met15Δ0 ura3Δ0 gcr1::KANMX ppt1::p616-YFP-Gcr1_bs-pPPT1-mCherry::NATMX6* |
| FW6415 | *MATa his3Δ1 leu2Δ0 met15Δ0 ura3Δ0 abf1::ABF1-V5-IAA7::KANMX6 leu2::pGPD1-osTIR1::LEU2 ppt1::p619-YFP-Abf1_bs-pPPT1-mCherry::NATMX6* |

| | |
|---|---|
| FW6411 | *MATa his3Δ1 leu2Δ0 met15Δ0 ura3Δ0*<br>*reb1::REB1-V5-IAA7::KANMX6 leu2::pGPD1-osTIR1::LEU2*<br>*ppt1::p620-YFP-Reb1_bs-pPPT1-mCherry::NATMX6* |
| FW6407 | *MATa his3Δ1 leu2Δ0 met15Δ0 ura3Δ0*<br>*ppt1::p592-YFP-pPPT1-mCherry::NATMX6* |
| FW629 | *MATa his3Δ1 leu2Δ0 met15Δ0 ura3Δ0* |
| FW8477 | *MATa his3Δ1 leu2Δ0 met15Δ0 ura3Δ0*<br>*rap1::RAP1-V5-IAA7::KANMX6 his3::pGPD1-osTIR1::HIS3*<br>*ura3::p375::URA3* |
| FW8523 | *MATa his3Δ1 leu2Δ0 met15Δ0 ura3Δ0*<br>*rap1::RAP1-V5-IAA7::KANMX6 his3::pGPD1-osTIR1::HIS3*<br>*ura3::p704::pTDH3-dCas9-3xFLAG-ADH1term::URA3*<br>*p692::pRS305-pSNR52-sgIRT2* |
| FW8531 | *MATa his3Δ1 leu2Δ0 met15Δ0 ura3Δ0*<br>*rap1::RAP1-V5-IAA7::KANMX6 his3::pGPD1-osTIR1::HIS3*<br>*ura3::p703::pTDH3-dCas9-Mxi1-3xFLAG-ADH1term::URA3*<br>*p692::pRS305-pSNR52-sgIRT2* |
| FW8527 | *MATa his3Δ1 leu2Δ0 met15Δ0 ura3Δ0*<br>*rap1::RAP1-V5-IAA7::KANMX6 his3::pGPD1-osTIR1::HIS3*<br>*ura3::p703::pTDH3-dCas9-3xFLAG-ADH1term::URA3*<br>*p705::pRS305-pSNR52-sgTEFI* |
| FW8535 | *MATa his3Δ1 leu2Δ0 met15Δ0 ura3Δ0*<br>*rap1::RAP1-V5-IAA7::KANMX6 his3::pGPD1-osTIR1::HIS3*<br>*ura3::p703::pTDH3-dCas9-Mxi1-3xFLAG-ADH1term::URA3*<br>*p352::pRS305-pSNR52-sgTEF1* |
| FW8529 | *MATa his3Δ1 leu2Δ0 met15Δ0 ura3Δ0*<br>*rap1::RAP1-V5-IAA7::KANMX6 his3::pGPD1-osTIR1::HIS3*<br>*ura3::p703::pTDH3-dCas9-Mxi1-3xFLAG-ADH1term::URA3*<br>*p706::pRS305-pSNR52-sgMLP1* |
