## Supplementary material for "RSC and GRFs confer promoter directionality by limiting divergent noncoding transcription": Table S3

**Table S4. Plasmids used in this study.**

| plasmid number | plasmid name |
|---|---|
| 592 | *YFP-pPPT1-mCherry::NATMX6* |
| 615 | *YFP-Cbf1_bs-pPPT1-mCherry::NATMX6* |
| 612 | *YFP-Gcn4_bs-pPPT1-mCherry::NATMX6* |
| 613 | *YFP-Cat8_bs-pPPT1-mCherry::NATMX6* |
| 614 | *YFP-Gal4_bs-pPPT1-mCherry::NATMX6* |
| 616 | *YFP-Gcr1_bs-pPPT1-mCherry::NATMX6* |
| 619 | *YFP-Abf1_bs-pPPT1-mCherry::NATMX6* |
| 620 | *YFP-Reb1_bs-pPPT1-mCherry::NATMX6* |
| 247 | *pNH605 pGPD1-osTIR1 LEU2* |
| 250 | *pNH603 pGPD1-osTIR1 HIS3* |
| 227 | *pWG444 (plasmid for making NAT deletion), NAT marker* |
| 627 | *3xFLAG-NATMX* |
| 704 | *pTDH3-dCas9-3xFLAG-ADH1term* |
| 703 | *pTDH3-dCas9-Mxi1-3xFLAG-ADH1term* |
| 692 | *pRS305-pSNR52-sgIRT2* |
| 706 | *pRS305-pSNR52-sgMLP1* |
| 352 | *pRS305-pSNR52-sgTEF1* |
| 252 | *pFA6A-V5-IAA7::KANMX6* |
