## Supplementary material for "RSC and GRFs confer promoter directionality by limiting divergent noncoding transcription": Table S4

**Table S5**. Oligonucleotide sequences used in this study.

| | | |
|---|---|---|
| IME1_NB_probe_R | ATGCAACGCCTACTTGTTTT | *IRT2* northern blot probe |
| IME1_NB_probe_F | GATGGAGGGTTGGCATAAAA | *IRT2* northern blot probe |
| SNR190_NB_probe_F | GGCCCTGATGATAATG | *SNR190* northern blot probe |
| SNR190_NB_probe_R | GGCTCAGATCTGCATG | *SNR190* northern blot probe |
| IRT2_RT_F | GACATCCGCATTCTTGCAGC | *IRT2* for RT-qPCR |
| IRT2_RT_R | CATGCTGTTCTTTCCGCCAC | *IRT2* for RT-qPCR |
| ACT1_RT_F | GTACCACCATGTTCCCAGGTATT | *ACT1* RT-qPCR |
| ACT1_RT_R | AGATGGACCACTTTCGTCGT | *ACT1* RT-qPCR |
| IRT2_gRNA_F | **ATGATATGTAG**TTTTAGAGCTAGAAATAGCAAGTTAAA | *IRT2* gRNA primer for cloning, highlighted is the gRNA sequence used |
| IRT2_gRNA_R | **TTAGATTATT**GATCATTTATCTTTCACTGCGGA | *IRT2* gRNA primer for cloning, highlighted is the gRNA sequence used |
| MLP1_gRNA_F | **GAGGCACGGAG**TTTTAGAGCTAGAAATAGCAAGTTAAA | *MLP1* gRNA primer for cloning, highlighted is the gRNA sequence used |
| MLP1_gRNA_R | **TAATAGATTT**GATCATTTATCTTTCACTGCGGA | *MLP1* gRNA primer for cloning, highlighted is the gRNA sequence used |
| Reb1bs_top | TCGGGTAACTTATCGGGTAACAT | Oligo used for cloning Reb1 binding site |
| Reb1bs_bottom | ATGTTACCCGATAAGTTACCCGA | Oligo used for cloning Reb1 binding site |
| Abf1bs_top | TTATCACTTCCCACGATTATCACTTCCCACGATT | Oligo used for cloning Abf1 binding site |
| Abf1bs_bottom | AATCGTGGGAAGTGATAATCGTGGGAAGTGATAA | Oligo used for cloning Abf1 binding site |
| Gal4bs_top | ACGGATTAGAAGCCGCCGAGCGGGCGACAGCCCTCCGACGGAAGACTCTCCTCCGT | Oligo used for cloning Gal4 binding site |
| Gal4bs_bottom | ACGGAGGAGAGTCTTCCGTCGGAGGGCTGTCGCCCGCTCGGCGGCTTCTAATCCGT | Oligo used for cloning Gal4 binding site |

| | | |
|---|---|---|
| Gcn4bs_top | TATGACTCATTCTATGACTCATTC | Oligo used for cloning Gcn4 binding site |
| Gcn4bs_bottom | GAATGAGTCATAGAATGAGTCATA | Oligo used for cloning Gcn4 binding site |
| Cbf1bs_top | T CACGTGA TCACGTGA T CACGTGA | Oligo used for cloning Cbf1 binding site |
| Cbf1bs_bottom | TCACGTG ATCACGTGA TCACGTG A | Oligo used for cloning Cbf1 binding site |
| Gcr1bs_top | TGGAAGCCTTGGAAGCCTTGGAAGCCT | Oligo used for cloning Gcr1 binding site |
| Gcr1bs_bottom | AGGCTTCCAAGGCTTCCAAGGCTTCCA | Oligo used for cloning Gcr1 binding site |
| Cat8bs_top | CCTTTAAGCCGACCTTTAAGCCGACCTT TAAGCCGA | Oligo used for cloning Cat8 binding site |
| Cat8bs_bottom | TCGGCTTAAAGGTCGGCTTAAAGGTCG GCTTAAAGG | Oligo used for cloning Cat8 binding site |
